# The regulatory landscape of the yeast phosphoproteome

**DOI:** 10.1101/2022.10.23.513432

**Authors:** Mario Leutert, Anthony S. Barente, Noelle K. Fukuda, Ricard A. Rodriguez-Mias, Judit Villén

## Abstract

The cellular ability to react to environmental fluctuations depends on signaling networks that are controlled by the dynamic activities of kinases and phosphatases. To gain insight into these stress-responsive phosphorylation networks, we generated a quantitative mass spectrometry-based atlas of early phosphoproteomic responses in *Saccharomyces cerevisiae* exposed to 101 environmental and chemical perturbations. We report phosphosites on 59% of the yeast proteome, with 18% of the proteome harboring a phosphosite that is regulated within 5 minutes of stress exposure. We identify shared and perturbation-specific stress response programs, uncover dephosphorylation as an integral early event, and dissect the interconnected regulatory landscape of kinase-substrate networks, as we exemplify with TOR signaling. We further reveal functional organization principles of the stress-responsive phosphoproteome based on phosphorylation site motifs, kinase activities, subcellular localizations, shared functions, and pathway intersections. This information-rich map of 25,000 regulated phosphosites advances our understanding of signaling networks.

**Highlights:**

- Ultra-deep reference yeast phosphoproteome covers 36,000 phosphorylation sites and reveals general principles of eukaryotic protein phosphorylation.
- High-dimensional quantitative atlas of early phosphoproteomic responses of yeast across 101 environmental and chemical perturbations identifies 25,000 regulated perturbation-phosphosite pairs.
- Identification of shared and perturbation-specific stress response phosphorylation programs reveals the importance of dephosphorylation as an early stress response.
- Dissection of the TOR signaling network uncovers subnetworks with differential stress responsiveness and points of pathway cross-talk
- Identification of functional organization of the phosphoproteome by dimensionality reduction and co-regulation analysis.

## Introduction

Cellular survival depends on conserved molecular mechanisms that maintain homeostasis under stress conditions. In the short term, these stress responses minimize acute damage to cell integrity and, in the long term, they promote adaptation and resilience. Stress response programs are orchestrated by complex signal transduction systems that sense environmental fluctuations and transmit information to various effector molecules. Their dynamics are mediated by signaling networks that rely on rapid and transient activities of kinases and phosphatases to coordinate cellular functions through reversible protein phosphorylation. Due to its broad impact on cellular function, phosphorylation-dependent signaling needs to be tightly controlled and aberrant signaling is involved in many diseases, including cancer.

The unicellular yeast S*accharomyces cerevisiae* harbors many conserved signaling pathways, encoding 159 protein kinases and phosphatases, 136 of which have human homologs, and therefore represents a valuable model organism to study phosphorylation networks at a systems level. Numerous studies have characterized environmental stress responses in yeast, exploring gene expression programs (Causton et al., 2001; Costanzo et al., 2021; Gasch et al., 2000; Hohmann and Mager, 2007), signaling pathways (Bahn et al., 2007; Gutin et al., 2015) and metabolic (Oliveira et al., 2015; Schulz et al., 2014), proteomic (Gutin et al., 2019; Paulo et al., 2015), and phenotypic responses (Brauer et al., 2008; Hillenmeyer et al., 2008; Viéitez et al., 2022). Several studies have employed phosphoproteomics to elucidate phosphorylation responses to individual perturbations (Gruhler et al., 2005; Kanshin et al., 2015a, 2015b; Lanz et al., 2021; Leutert et al., 2019; MacGilvray et al., 2018; Oliveira et al., 2012, 2015; Saleem et al., 2010; Smolka et al., 2007; Vaga et al., 2014) or to the activation of specific pathways (Dokládal et al., 2021; Holt et al., 2009; Plank et al., 2020; Soste et al., 2014). Systems-level genetic perturbation studies have applied phosphoproteomics (Bodenmiller et al., 2010; Li et al., 2019) or orthogonal measurements (Schulz et al., 2014; da Silveira Dos Santos et al., 2014; Viéitez et al., 2022; van Wageningen et al., 2010) to uncover kinase-substrate relationships and functional phosphosites in yeast. While these studies have made substantial progress in our understanding of kinases and phosphosites, a systematic view of the activation states and architecture of the stress-responsive phosphorylation signaling network is still missing.

To address this gap we profiled quantitative phosphoproteomic responses in over 600 samples exposed to 101 environmental and chemical perturbations to provide one of the largest and most systematic perturbation-phosphoproteomic resources for any species. This unprecedented scaling of phosphoproteomic experiments to hundreds of samples without compromising identification depth or quantitative measurement precision was enabled by combining our recently developed high-throughput sample preparation strategies, mass spectrometry acquisition methods, and computational approaches (Lawrence et al., 2016; Leutert et al., 2019; Searle et al., 2018, 2019). The depth and high-dimensionality of our integrated resource allowed us to uncover fundamental properties of protein phosphorylation, identify shared and perturbation-specific stress response programs, reveal the extent of dephosphorylation, dissect the TOR signaling network and uncover general principles that underlie functional phosphoproteome organization.

## Results

### The phosphoproteomic landscape of *S. cerevisiae* exposed to 101 cellular perturbations

To study the regulatory and functional properties of protein phosphorylation, we studied phosphoproteomic responses of haploid yeast, exposed to 101 diverse perturbations (Figure 1A). These perturbations covered broad classes of environmental and chemical conditions as well as drug treatments and systematically targeted diverse aspects of yeast cell biology (Figure 1B, Table S1). For the majority of perturbations, yeast was treated for 5 minutes, in order to capture early signaling responses. We reasoned that functional phosphorylation of specific substrates occurs rapidly following stimulation, while promiscuous phosphorylation and changes in transcript and protein abundance takes place more slowly (Gasch et al., 2000; Kanshin et al., 2015a). We generated an ultra-deep reference yeast phosphoproteome from a pooled sample of all 5-minute perturbations by multi-protease digestion, deep offline fractionations using two different peptide separation methods, and data-dependent acquisition (DDA) MS. The number of phosphosite identifications obtained with trypsin plateaued at ~30,000 phosphosites and use of alternative proteases provided an additional ~6,000 phosphosites (Figure 1D). Collectively, this resulted in 36,405 high-confidence phosphosites at a peptide-spectrum match false discovery rate (FDR) of <1% and localization probabilities of > 95% (Ascore >13) (Figure 1C, Table S2) (Beausoleil et al., 2006). Of the identified phosphosites, 40% (14,271 phosphosites) have not been annotated in the Saccharomyces Genome Database (SGD, https://www.yeastgenome.org/) (Figure 1E). Many of these novel phosphosites were derived from non-tryptic peptides (30%) and/or are of low abundance (80% of novel phosphosites were identified by a single spectrum); others might be perturbation-specific. Phosphosites that were present in the SGD but not identified here (Figure 1E) potentially occur under different growth conditions, or originate from differences in phosphosite localization and filtering criteria or inadequate aggregation of proteomic data leading to substantial accumulation of false positives (Lanz et al., 2021; Ochoa et al., 2019).

**Figure 1.**
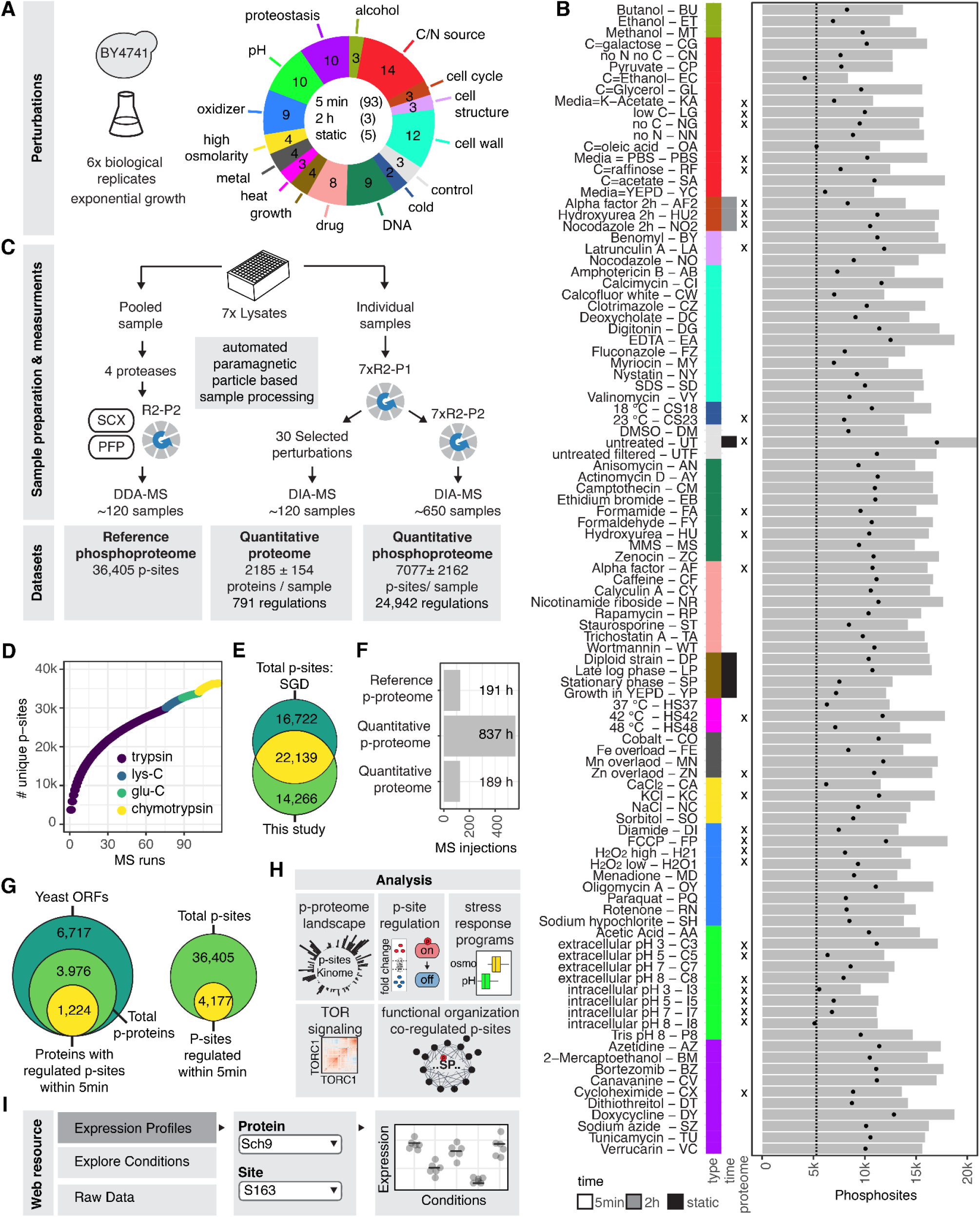
Mass spectrometry-based atlas of phosphoproteomic responses to 101 cellular perturbations. **(A)** Overview of broad perturbation types and numbers and timing of all treatments. The color scheme for the perturbation types is consistent across all figures. **(B)** Description of all treatments and indication if total proteome was measured (marked with x). The bar plot shows total quantified phosphosites per treatment. Black dots indicate the number of phosphosites measured in at least two replicates and the dashed line indicates the number of core phosphosites. C = carbon source, N = nitrogen source, Media = media exchange. **(C)** Schematic representation of sample preparation workflows, MS measurements, created datasets and key numbers. **(D)** Cumulative identification of unique phosphosites across MS measurements within the reference phosphoproteome. **(E)** Overlap of our reference phosphoproteome with all phosphosites reported by SGD. **(F)** Number of MS injections and total effective MS acquisition time for each dataset. **(G)** Left: count of all reported open reading frames (ORF) by SGD, proteins identified in our reference phosphoproteome and proteins with quantitatively regulated phosphosites within 5 min. Right: overlap of the total number of identified phosphosites and phosphosites regulated within 5 min. **(H)** Overview of performed analyses. **(I)** Overview of web resource for interactive data exploration (https://yeastphosphoatlas.gs.washington.edu).

### Quantitative atlas of phosphoproteomic stress responses

To quantitatively measure the phosphoproteomes in response to all 101 perturbations, we prepared phosphoproteomic samples using our recently developed R2-P2 workflow (Leutert et al., 2019) and analyzed them by data-independent acquisition (DIA) MS. To control for technical variation, we performed block randomization across the whole experiment and included sample preparation and MS measurement quality control samples at every stage (Figure S1A). We further measured the total proteomes from matching samples for 30 selected perturbations, covering all broad perturbation types. The data was filtered to a global 1% precursor FDR and phosphosite localization probabilities of >75% (Figure S1B). Our approach enabled deep phosphoproteome coverage and robust quantitation across all samples (Figure S1C-S1E). On average, we identified and individually quantified 7,077 ± 2,162 (mean and standard deviation) phosphosites per phosphoproteome sample (Table S3, Figure 1B and 1C) and 2,185 ± 154 proteins per total proteome sample (Table S4, Figure 1C). The final datasets encompassed 811 mass spectrometry injections, corresponding to 51 days of effective data acquisition time (Figure 1F).

A main challenge for multi-condition phosphoproteomic experiments are missing quantifications due to low phosphopeptide abundance or ambiguous phosphosite localization. We applied a stringent threshold requiring that a localized phosphosite be quantified in at least 50% of all samples, which provided a good balance between quantitative reproducibility and retained phosphosites (Figure S2A and S2B). Using this filtering approach, we obtained robust quantification of 5,284 phosphosites with an overall coverage of 66.5% (Figure S2C). We refer to this set as the core phosphoproteome (Figure 1B). We performed imputation assuming missing data is due to low intensity values (Figure S2D) and showed that technical variance was largely explained by separately processed 96-well plates (Figure S2E and S2F). We were able to correct the data for sample preparation batch effects, so that the treatments explained 35% of the variance (Figure S2G-S2J). Our approach resulted in a complementary pair of core phosphoproteome datasets: an uncorrected dataset (Table S3) that was used for linear modeling of differential phosphosite abundance and a corrected dataset (Table S5) that was used to compare phosphosite abundances across the whole dataset and to perform co-regulation analysis.

To understand how individual perturbations affect relative phosphorylation of sites across the proteome, we performed linear modeling to test all phosphosite-treatment pairs (529,400 total tests) for differential abundance against the untreated samples (Fig S3A, Table S6). Considering all 5-min perturbations, a >2-fold change and significance at an adjusted p-value of <0.05 resulted in 24,942 regulated perturbation-phosphosite pairs distributed across 4,177 unique phosphosites on 1,155 proteins (Figure 1C).

In summary, we identified 36,405 phosphosites on 3,976 proteins, which account for 59% of all annotated open reading frames in *S. cerevisiae* (Figure 1G). At least 18% of all proteins (1,224 proteins) contain a phosphosite that was significantly regulated within 5 min of perturbation (Figure 1G). This translates to 11% (4,177) of all identified phosphosites showing regulation under at least one perturbation. We integrated these high-dimensional datasets in multiple ways that showcase how this resource can used to learn about the organization of signaling systems, and provide the underlying data for further exploration at https://yeastphosphoatlas.gs.washington.edu (Figure 1H and 1I).

### Characteristics of early stress-responsive phosphosites

We performed comparative analysis of the global and the regulated phosphoproteome to identify sequence, structural and functional characteristics of early stress-responsive phosphosites. We found no bias for protein abundance for single-, multi-, or hyperphosphorylated proteins within the reference phosphoproteome, whereas the regulated phosphoproteome showed a slight shift towards more abundant proteins. This can be explained by the reduced coverage of the core phosphoproteome together with the data completeness filter which are biased towards higher abundant peptides (Figure 2A).

**Figure 2.**
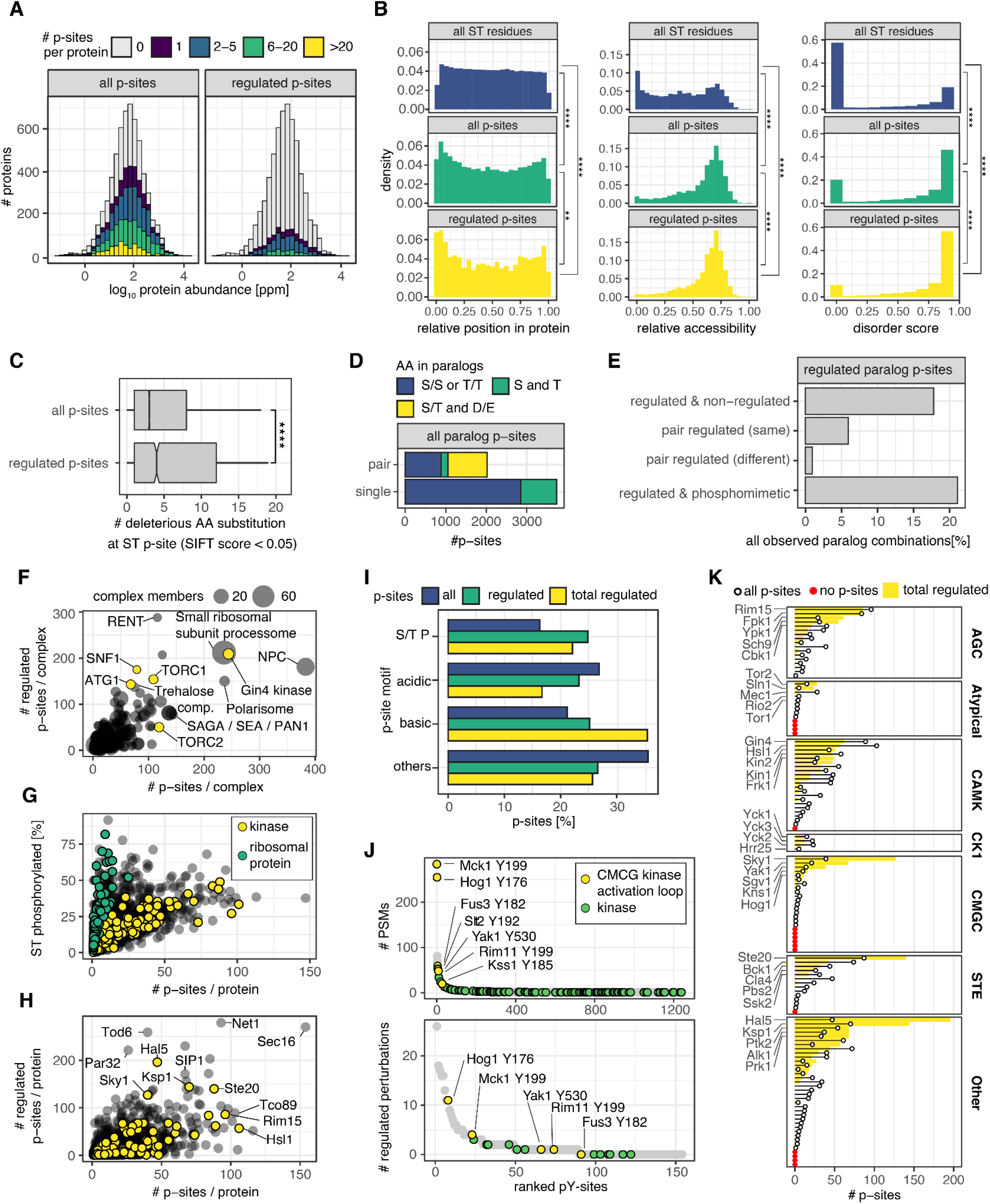
Structural, regulatory, and functional features of the yeast phosphoproteome. **(A)** Phosphosite count per protein mapped on protein abundance distribution for identified and regulated phosphosites. **(B)** Density plots depicting the relative residue position within a protein sequence (left column), the relative solvent accessibility of a residue within the protein structure (middle column) and a prediction score for the residue to be in a disordered protein region (right column) for all S and T residues within all phosphoproteins (top row), all identified phosphosites (middle row) and all regulated phosphosites (bottom row). KS test adjusted p-value: ** < 0.01, *** < 0.0001. **(C)** Count of amino acid substitutions with a SIFT score < 0.05 for all phosphosites and regulated phosphosites. Wilcoxon test p-value: **** p < 0.0001. **(D)** Comparison of phosphosites on paralog proteins where both paralog proteins (pair) are modified at the same conserved site (S/S or T/T), are modified at the same position but different amino acid (S and T), one of the paralog pairs shows a phosphomimetic amino acid substitution at the same site (S/T and D/E) and cases where only one paralog is modified but the other paralog has a T or S at the same side (single). **(E)** Comparison of regulated phosphosites on paralogs that have a measured phosphosite at the same conserved site or one of the paralogs has a phosphomimetic substitution. **(F)** Regulated phosphosite-perturbation pairs versus all phosphosites identified for protein complexes. Kinase containing complexes are marked in yellow. **(G)** Percentage of phosphorylated S and T versus number of phosphosites on the same protein. **(H)** Number of regulated phosphosite-perturbation pairs for a protein versus all phosphosites identified on the same protein. **(I)** Percentage of phosphosites associated with a specific motif type for all identified phosphosites, regulated phosphosites and regulated phosphosite-perturbation pairs. **(J)** All identified pY sites ranked by number of peptide spectrum matches (top) and pY sites ranked by number of conditions that lead to regulation of the phosphosite (bottom). **(K)** All identified phosphosites and regulated phosphosite-perturbation pairs across the entire yeast kinome grouped by kinase class. The top 5 kinases with the most regulated perturbations per class are annotated. Kinases without any detected phosphosites are marked in red.

Next, we compared unmodified Ser/Thr residues in identified phosphoproteins versus all phosphosites and all regulated phosphosites in terms of their relative position in protein sequences, relative accessibility, and occurrence within disordered protein regions (Figure 2B) (Jumper et al., 2021; Pentony et al., 2010). Interestingly, we found strong enrichment of phosphorylation occurring on protein N-termini and, to a lesser extent on C-termini. This effect was even more pronounced for regulated phosphosites (Figure 2B). Enrichment in C-terminal phosphorylation has been described before in mammals (Villén et al., 2007), whereas N-terminal phosphorylation is less defined, but might be involved in co-translational regulation. Phosphosites in general, especially regulated phosphosites, were also enriched in accessible protein regions and in disordered protein regions (Figure 2B).

To assess the functional relevance of phosphosites, we extracted ‘sorting intolerant from tolerant substitutions’ (SIFT) scores from the mutfunc database (Ng and Henikoff, 2001; Wagih et al., 2018). SIFT scores are based on sequence homology and physical properties of amino acids and predict whether an amino acid substitution affects protein function. We found that residues harboring regulated phosphosites had a higher number of amino acid substitutions predicted to be deleterious by SIFT (Figure 2C). This result suggests that regulated phosphosites are more likely to occur at regions of high sequence conservation and impact protein function (Studer et al., 2016).

To further investigate phosphosite conservation we compared yeast protein paralogs, which arose by whole-genome or single-gene duplication (Byrne and Wolfe, 2005). We only considered phosphosites on paralog pairs that were located in paralog-specific peptides and where the modified residue was either conserved in both paralogs; mutated from Ser to Thr; or phosphorylated in one paralog and a phosphomimetic Asp or Glu residue in the other paralog. After applying these criteria, we were left with 5,754 paralog site-pairs (across 722 paralog-pairs). We find that for a third of paralog phosphosite-pairs, phosphorylation is conserved or complemented with a phosphomimetic mutation (Figure 2D). In most of these cases, only one paralog site is regulated across perturbations, suggesting that differential phosphorylation can contribute to diversification of paralog function across environmental conditions (Figure 2E). Strikingly, we also found a number of paralog pairs (13%) where a regulated phosphosite in one paralog is substituted by a phosphomimetic residue in the other paralog. Such an evolutionary switch might lead to functional diversification where one paralog is constitutively active, while the other paralog has a phosphorylation-dependent condition-specific function.

Next, we investigated which proteins and protein complexes are highly phosphorylated. We found that the nuclear pore complex (NPC) harbors the most phosphosites (383 phosphosites) with 181 regulated phosphosite-perturbation pairs (Figure 2F). The NPC is known to undergo cell cycle dependent phosphorylation (Aitchison and Rout, 2012; Holt et al., 2009) and the observed phosphoregulation is likely a consequence of stress-induced cell cycle changes. Kinases were the protein class with the most phosphosites per protein and ribosomal proteins showed the highest percentage of Ser and Thr that are phosphorylated (Figure 2G). Remarkably, several proteins showed >100 regulated phosphosites (Figure 2H).

Overall, our analyses revealed that regulated phosphosites occur more frequently on protein termini and in accessible or disordered regions and have a higher probability of conservation and impact protein function.

### Regulated phosphosites of the yeast kinome

We assessed the distribution of broad phosphosite motif classes across the reference and regulated phosphoproteome (Mok et al., 2010; Villén et al., 2007). We found that ~15% of all phosphosites occurred in proline-directed motifs (targeted by members of the CMCG kinase class, which included MAPKs, CDKs and Yak1), ~25% in acidic motifs (targeted by CKs, GSK3 homologs Mrk1 and Rim11 and others) and ~20% in basic motifs (targeted by >35 basophilic kinases most prominently in the AGC and CaMK kinase groups, which include PKA, Snf1 and Sch9) (Figure 2I). Regulated phosphosites occurred more often in proline-directed and basic motifs, accounting for >50% sites and less often in acidic motifs. Phosphorylation on acidic motifs tends to have higher stoichiometry (Wu et al., 2011) and our results suggest they represent constitutively active instead of stress-sensitive phosphorylation. (Figure 2I). Strikingly, when considering all regulated phosphosite-perturbation pairs, we found that 35% of regulated phosphosites have a basic motif (Figure 2I).

Tyrosine phosphorylation is thought to be low in yeast and dependent on a few dual-specificity kinases (Brinkworth et al., 2006). We found 1,243 phosphotyrosine sites, although the majority of these (64%) were identified with only one spectrum match, indicating that they are either low abundant or mislocalized (Figure 2J). Of all tyrosine phosphorylation 10% occurred on kinases. Highly-abundant phosphotyrosine sites (identified with >20 spectral matches) included 7 instances of CMCG kinase activation loop phosphorylation (Figure 2J). In the core phosphoproteome, we found only ~90 phosphotyrosine sites that were regulated in at least one perturbation (Figure 2J). Taken together, our large-scale study confirms low levels of tyrosine phosphorylation in yeast with increased occurrence on kinases.

Mapping phosphosites across the kinome emphasized overall high numbers of modified and regulated residues, particularly for AGC, CAMK and STE kinase families (Figure 2K). CMGC kinases mediate response to diverse signals and we find abundant phosphorylation on stress responsive Sky1, Yak1 and Hog1 kinases. Only for 6 CMGC kinases, we detected no phosphosites. Three of these, Smk1, Kdx1 and Ime2, are directly involved in sporulation and therefore unlikely to be activated under the tested conditions. Other kinases with low numbers of phosphosites such as Tor1, Tor2, and Snf1 are active within complexes and we find very high numbers of regulated phosphosites on their associated subunits (Tco89 and Kog1 for TORC1/2, and Sip1 and Gal83 for Snf1 complex). Overall, only 16 out of 129 predicted yeast kinases had no detected phosphosites.

### Dissecting stress responsive phosphorylation sites

Differential phosphosite regulation across perturbations can show the extent of the global signaling response for each perturbation and give insights into the function of individual phosphosites. We analyzed core phosphosites that are significantly regulated (>2-fold change and adjusted p-value < 0.05) for each perturbation compared to untreated yeast (Figure 3A). Phosphoproteome changes were more widespread upon pH perturbations, osmotic stresses, and nutrient perturbations; phosphosites were mostly down-regulated upon cell wall perturbations. Agents that affected proteostasis or DNA damage and most drug treatments caused overall only modest responses at core phosphosites (Figure 3A and 3B). Regulation of phosphosites within 5 minutes of perturbation could not be explained by changes in protein abundance. By analyzing the proteome of 30 selected perturbations we found that most perturbations regulated less than 3% of all measured proteins (Figure S3B-S3E) and that this regulation does not correlate with changes in phosphorylation (Figure 3C).

**Figure 3.**
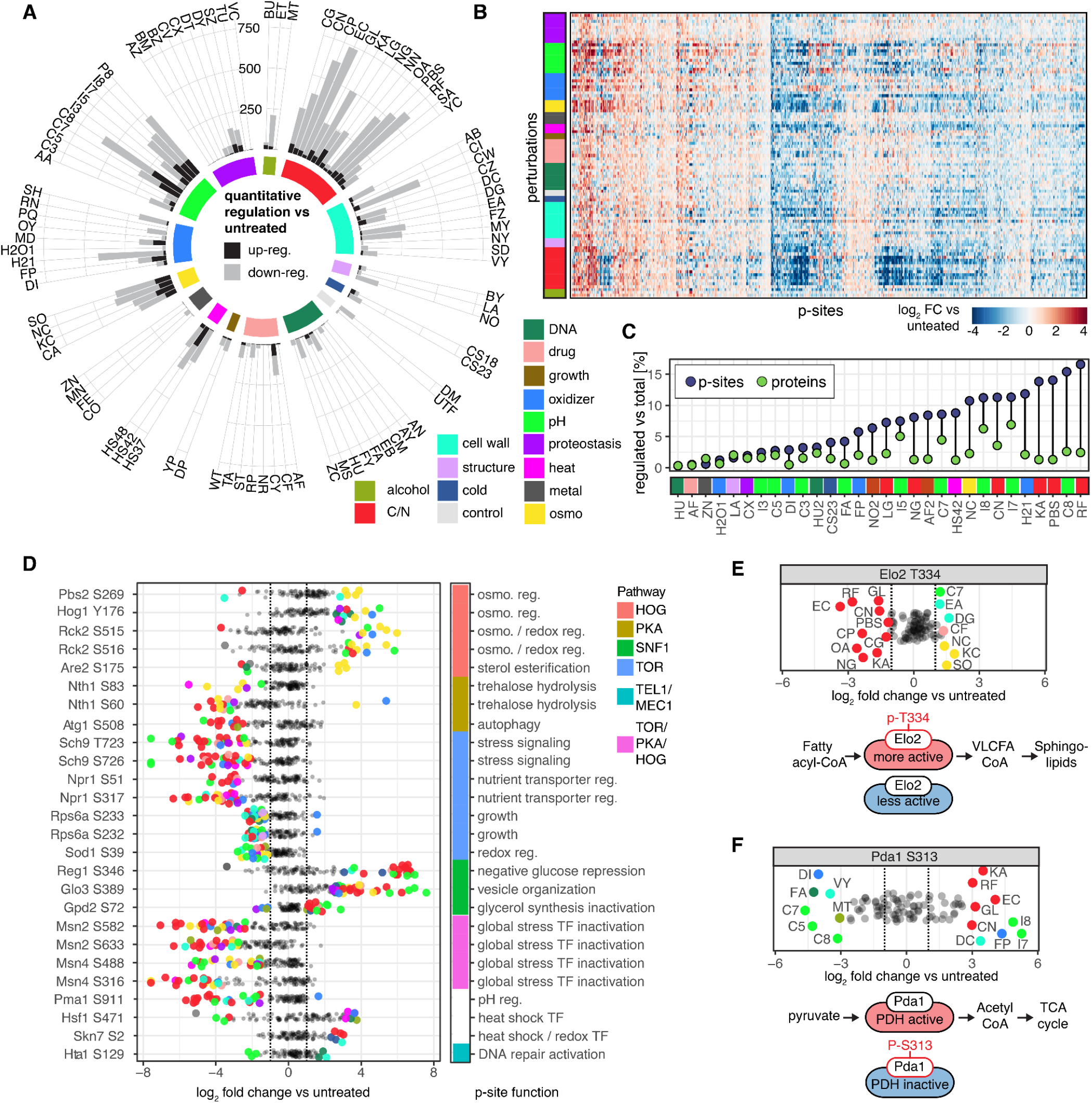
Dissecting stress responsive phosphorylation sites. **(A)** Up- and down-regulated phosphosites across all 5-min perturbations versus untreated control ordered by perturbation types. **(B)** Heatmap of log_2_ fold changes of individual phosphosites in all perturbations versus the untreated condition. Perturbation types are color coded as in (A). Unsupervised hierarchical clustering is performed on phosphosite log_2_ fold changes. Only phosphosites that are regulated in at least one perturbation are considered. **(C)** Percentage of proteins and phosphosites that are regulated within a perturbation compared to all proteins and phosphosites considered for the perturbation. Perturbations are color coded as in (A). **(D)** Log_2_ fold changes (versus untreated) of well-characterized phosphosites. Each dot represents a perturbation and is shown color coded as in (A) if the indicated phosphosite is significantly regulated by >2-fold (adjusted p-value <0.05). Pathway or upstream kinase for each phosphosite and the known function of the phosphosite is indicated. **(E)** Top: same as (D) for Elo2 T334 phosphorylation. Bottom: Elo2 acts on fatty acids and phosphorylation of T334 increases its activity, leading to increased amounts of very long chain fatty acids (VLCFA), resulting in increased sphingolipid synthesis. **(F)** Top: same as (D) for Pda1 S313. Bottom: Pda1 S313 phosphorylation inactivates the pyruvate dehydrogenase (PDH) complex and inhibits conversion of pyruvate to acetyl-CoA, thereby blocking cellular respiration and favoring fermentation.

Next, we focused on individual phosphosites with well-characterized functions and investigated the regulation of these sites across all tested perturbations (Figure 3D, Figure S3F). We highlight here some phosphosites and describe additional examples in the Supplementary Text. First we analyzed activating phosphosites within the high osmolarity glycerol (HOG) MAP kinase pathway, which is well studied for its essential functions in the osmotic stress response (Romanov et al., 2017). In the canonical HOG pathway the MAP kinase kinase Pbs2 is activated through phosphorylation by an upstream MAP kinase kinase kinase and in turn leads to phosphorylation and activation of the MAP kinase Hog1. Hog1 phosphorylates multiple effector proteins, including Rck2 and Are2 with roles in cell cycle regulation and membrane maintenance, respectively. We found robust up-regulation of activating phosphosites on Pbs2, Hog1, Rck2 and Are2 across all four osmotic stresses (Figure 3D). Whereas Pbs2, Rck2 and Are2 activation was mostly specific for osmotic stresses, Hog1 was also activated in mitochondrial perturbations, heat shock, and DNA damage conditions. These distinct activation patterns in the HOG pathway suggest there are multiple ways of inducing Hog1 and additional regulatory layers to create downstream specificity. Hog1 activation independent of upstream MAP kinase kinases could be explained by stress-specific inhibition of protein tyrosine phosphatases (Ptp2 and Ptp3) that maintain Hog1 in an inactive state under normal conditions (Lee and Levin, 2018). Additionally, we identified many other regulated phosphosites on Pbs2, Rck2, and Are2 that may modulate specific activation states.

The transcription factors Msn2 and Msn4 are central mediators of the environmental stress response (ESR) gene expression program (Gasch et al., 2000), integrating multiple stress and nutrient sensitive pathways, including the HOG pathway. Msn2 and Msn4 activity is dependent on their complex and transient phosphorylation states. Short-lived multisite dephosphorylation of Msn2 and Msn4 in their nuclear localization signal (NLS) is associated with Msn2/4 translocation to the nucleus and induction of ESR transcription (De Wever et al., 2005; Reiter et al., 2013). In contrast to the specific regulation of HOG pathway effector proteins discussed above, we observe dephosphorylation of Msn2 and Msn4 phosphosites in their NLS for almost all broad perturbation types (Figure 3D). Interestingly, a wide quantitative range of dephosphorylation was observed for these phosphosites. We found strongest dephosphorylation in nutrient perturbations and lower dephosphorylation in osmotic, oxidative and proteostasis stresses. This magnitude difference is likely a consequence of pathway crosstalk and/or feedback leading to a fine-tuned regulation of ESR gene expression programs.

Strikingly, a few phosphosites showed bidirectional regulation in response to multiple perturbations. This is the case for key regulatory phosphosites on the two metabolic enzymes Elo2 and Pda1. We observed increased phosphorylation of the fatty acid elongase Elo2 at T334 upon osmotic stress, digitonin, EDTA, caffeine and pH and decreased phosphorylation under nutrient limitation and in the majority of carbon source changes (Figure 3E). Activating Elo2 phosphorylation at T334 has previously been shown to be required for modulation of sphingolipid synthesis (Viéitez et al., 2022; Zimmermann et al., 2013). Elo2 T334 phosphorylation increases sphingolipid production to potentially enhance membrane integrity upon perturbations that impact cell membrane structure, while T334 dephosphorylation upon starvation leads to decreased sphingolipid production. The regulation we observe suggests that Elo2 T334 acts as a rapid and broad stress responsive regulator of the sphingolipid biosynthesis pathway, with possible implications for metabolic fluxes, membrane dynamics and autophagy regulation. Bidirectional regulation was also found for the highly conserved, inactivating phosphorylation of the pyruvate dehydrogenase complex (PDH) subunit Pda1 on S313 that acts as a central metabolic switch between fermentation and respiration (Oliveira et al., 2012; Uhlinger et al., 1986) (Figure 3F and Supplementary Text).

In summary, we show that integration of the phosphorylation state of individual phosphosites across a large number of perturbations can be used to understand common and specific cellular responses to different perturbations, as exemplified here for phosphosites with well-characterized functions. We observed that many phosphosites with known functions responded in an expected manner to certain perturbations, but were also regulated by seemingly unrelated or unexpected perturbations. Interestingly, we find that several key functional phosphosites show large differences in fold changes across perturbations, which can be generated by feedforward loops (Goentoro et al., 2009). We believe that our resource can be used to identify potential new functions of known phosphosites and signaling network motifs that can be followed up in future experiments.

### The shared phosphorylation stress response program is defined by nutrient sensing pathways and broad dephosphorylation

We next examined general trends in regulation across the yeast phosphoproteome. We first assessed the regulatory behaviors of individual phosphosites across all perturbations, and found that most phosphosites are regulated unidirectionally (i.e. the phosphosite is either consistently up- or down-regulated compared to untreated yeast) (Figure 4A). When taking a perturbation-centric approach (i.e. assessing the regulatory behaviors of all phosphosites within a perturbation), we discovered that down-regulation of phosphorylation was more frequent compared to upregulation in nearly every perturbation (Figure 4B). This effect did not appear to be due to data processing and imputation. Interestingly, these results suggest a central role for dephosphorylation during the initial stress response. Indeed, phosphatases occupy key positions in the stress response network (Figure S4A) (Ariño et al., 2019) and phosphatase deletion strains have shown a high number of stress sensitive growth phenotypes (Figure S4B) (Viéitez et al., 2022). To investigate if dephosphorylation is a general feature of the early stress response in yeast we analyzed published temporal phosphoproteomic data (Kanshin et al., 2015b; Leutert et al., 2019; MacGilvray et al., 2018). We found that, for the majority of cases, down-regulation of phosphorylation outweighed up-regulation, specifically at early time points (Figure S4C. We also examined global phosphoproteomic responses to environmental stresses for other species (*C. elegans*, *H. sapiens*, *R. rattus*), and found a similarly-strong dephosphorylation response, suggesting that global dephosphorylation might be a conserved stress responses (Figure S4D and S4E) (Huang et al., 2020; Needham et al., 2019; Rigbolt et al., 2014).

**Figure 4.**
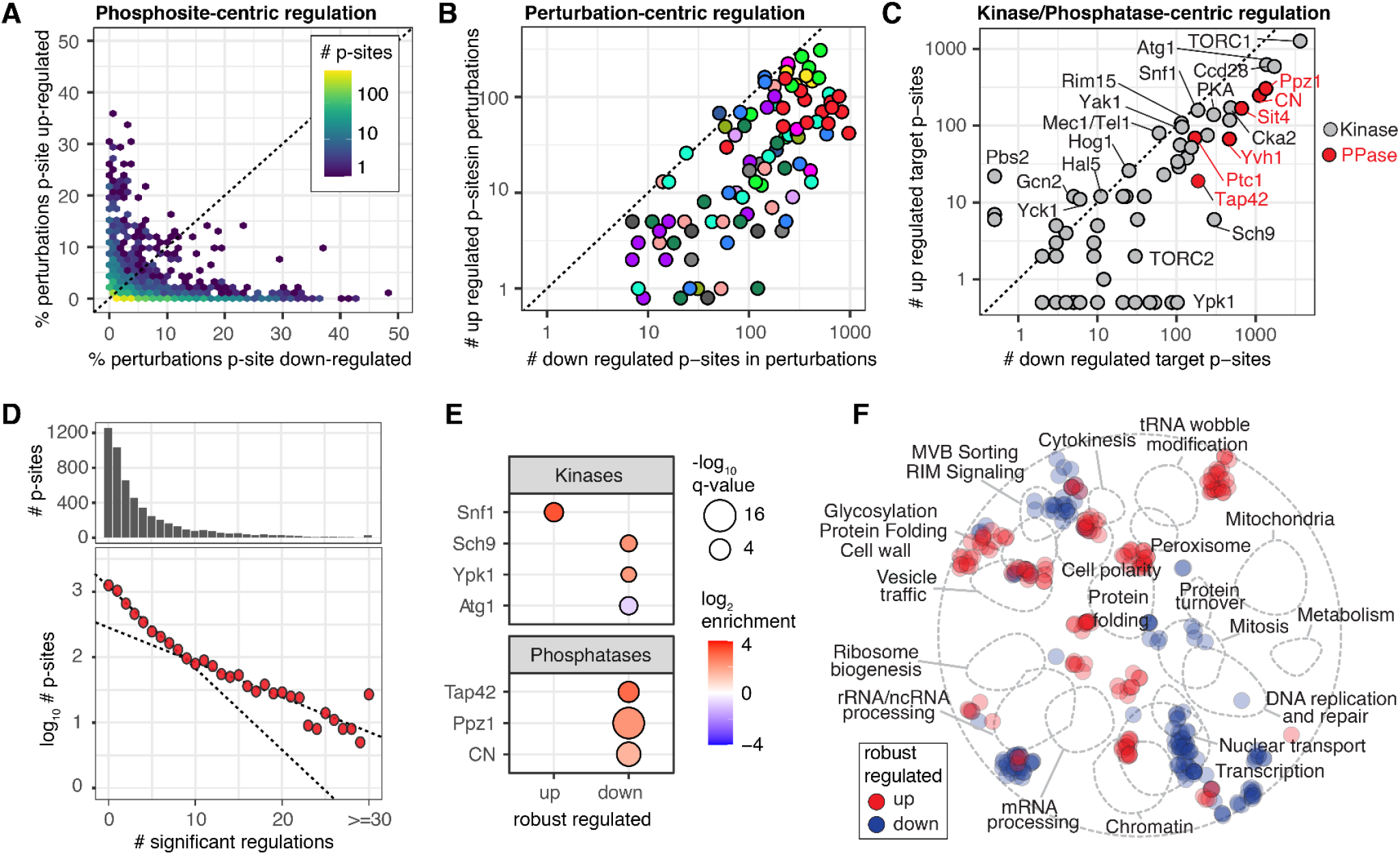
Shared phosphorylation stress response program. **(A)** Percentage of perturbations where a phosphosite was up-versus down-regulated. **(B)** Counts of up-versus down-regulated phosphosites for every perturbation represented in log_10_ scale. Perturbations are color coded according to Figure 1. **(C)** Counts of up-versus down-regulated phosphosites targeted by a specific kinase or phosphatase represented in log_10_ scale. **(D)** Top: Histogram of phosphosites with significant regulations. Bottom: same data, but plotting the y-axis in a log_10_ scale with linear fits showing a change in slope at 10 regulations (shared response). **(E)** Significantly enriched (Fisher exact test q-value < 0.01) kinase and phosphatase substrates for robustly regulated phosphosites. **(F)** Phosphoproteins that contained robustly up- or down-regulated phosphosites were mapped onto the global yeast genetic-interaction network and SAFE analysis was performed. Proteins in the network that are significantly enriched for functional interactions with phosphoproteins are indicated as color coded dots. Proximity to annotated subnetworks (dashed lines) indicates related cellular function or localization.

To dissect the impact of different kinases and phosphatases on the directionality of phosphosite regulation, we grouped phosphosites by reported upstream kinase or phosphatase and counted perturbations that led to up- or down-regulation (Figure 4C). This analysis revealed kinases that are strongly associated with up-regulated phosphosites such as Snf1, Rim15 and Yak1, all part of glucose sensing systems; Hog1, Pbs2 and Hal5, involved in osmo- and halotolerance; and Mec1/Tel1 involved in DNA damage and cell cycle control. Kinases and phosphatases that were associated with down-regulated phosphosites, included TOR and PKA pathway-related kinases (TORC1, Atg1, Sch9, Ypk1) and phosphatases (Tap42, Sit4), Cka2 kinase, the pH stress activated Ppz1 phosphatase, calcineurin phosphatases and the Ptc1 phosphatase (Ariño et al., 2019; González and Hall, 2017).

Next, we investigated whether a shared stress response program exists across the yeast phosphoproteome. We found that the number of perturbations where a phosphosite was significantly regulated decreased exponentially and we identified a breakpoint in the exponential distribution at around 10 perturbations, with a slower dropoff after this point (Figure 4D). Considering this, we defined phosphosites that are unidirectionally regulated in 10 or more perturbations as a shared phosphoproteomic stress response, and identified responsible kinases and phosphatases. We found that upregulated sites were enriched in Snf1 substrates whereas downregulated sites were enriched in targets of the Sch9 and Ypk1 kinases and Ppz1, calcineurin, and Tap42-PP2A/Sit4 phosphatases (Figure 4E). To gain insight into biological processes affected by the shared phosphorylation stress response, we mapped proteins containing at least one regulated phosphosite onto the yeast genetic-interaction network and performed Spatial Analysis of Functional Enrichment (SAFE) analysis (Baryshnikova, 2016; Usaj et al., 2017) to identify regions of the network that are significantly enriched for interactions with regulated phosphoproteins (Figure 4F). We found enriched functions for proteins with up-regulated phosphosites in processes associated with cell polarity and proteins with down-regulated phosphosites in nuclear processes such as transcription and nuclear transport as well as mitosis.

In summary, we found that deactivation of signaling pathways that are constitutively active under exponential growth conditions (TOR, PKA) and the action of stress responsive (calcineurin, Ppz1) and TOR associated (Tap42-PP2A/Sit4) phosphatases play a major role in shaping the phosphoproteome during the initial shared stress response. This signature is consistent with a shift from biosynthetic processes to catabolic processes. Unlocking phosphatase activity upon stress exposure might provide the cell with a fast mechanism to rewire growth signaling and homeostatic functions.

### Perturbation-specific phosphorylation response programs

The high dimensionality of our dataset also allowed us to identify novel regulatory connections between stress responses. We determined stress response similarities by hierarchical clustering of a correlation matrix (Figure 5A). Perturbations were nearly all uncorrelated or positively correlated, which suggests that, generally, when pathways are regulated they are regulated in the same way. As expected, a series of prominent clusters of known related perturbations are visible in the data, such as the osmotic stresses, pH stresses, and nutrient perturbations. Interestingly, this approach was able to further resolve nutrient perturbations into carbon source exchange and limitation.

**Figure 5.**
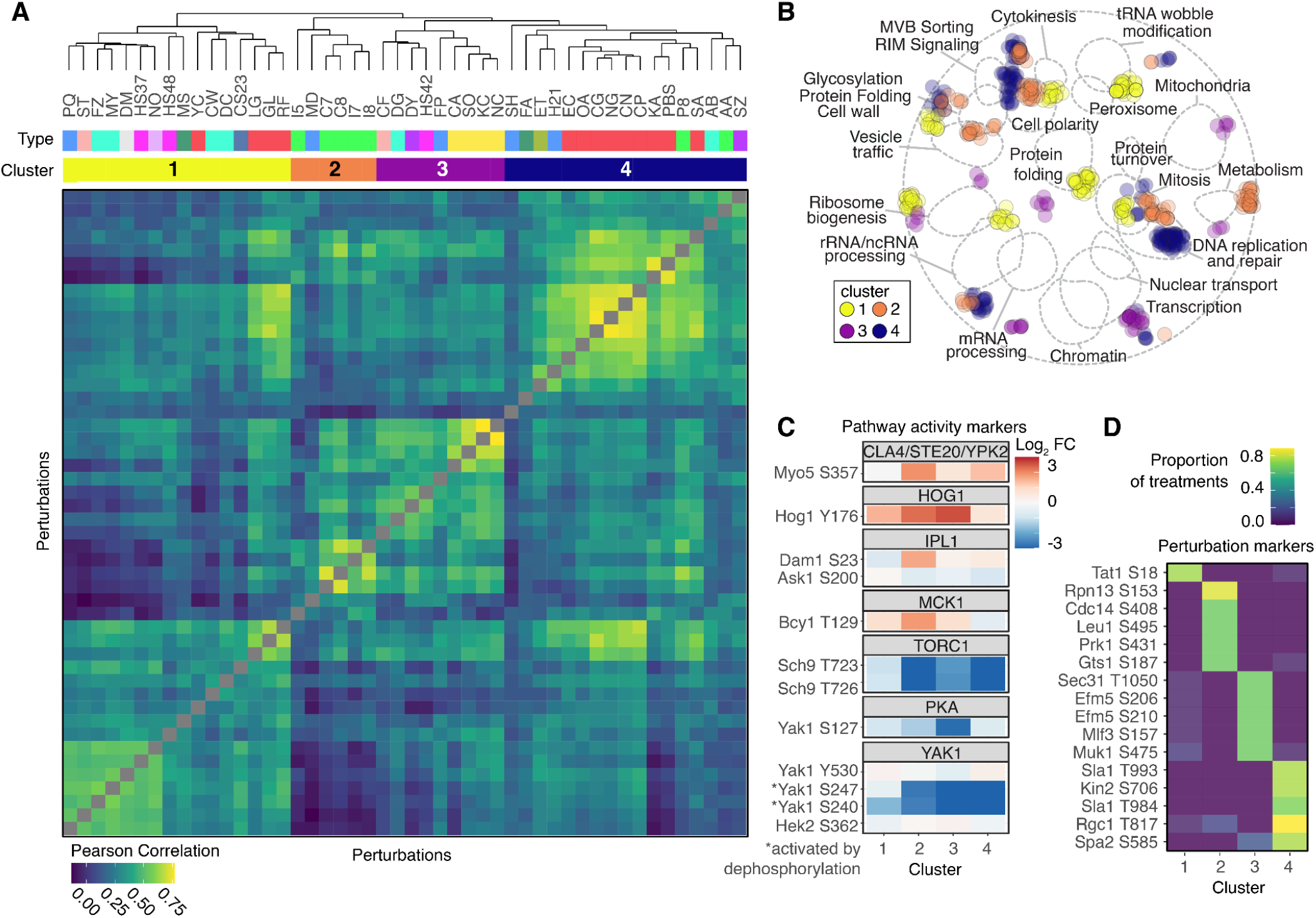
Perturbation-specific phosphorylation stress response programs. **(A)** Pearson correlation heatmap of phosphosite log_2_ fold changes (versus untreated) for selected perturbations. Perturbations that were selected if the number of regulated phosphosites was in the upper 50th percentile and contained phosphosites regulated in at least one other perturbation. Rows and columns were arranged by hierarchical clustering and the resulting dendrogram is displayed along with perturbation types. Four major perturbation clusters were determined by cutting the dendrogram at a uniform height. **(B)** Same as Figure 4F but for phosphoproteins that contain phosphosites specific to each of the 4 perturbation clusters. **(C)** Mean log_2_ fold change in each perturbation cluster for phosphosites that act as markers for pathway activity. **(D)** Representative highly-specific marker phosphosites for different perturbation clusters.

We find four main perturbation clusters, corresponding to major phosphorylation stress response programs that target proteins involved in functionally different biological processes (Table S7, Figure 5B). To obtain insight into the activity of signaling pathways across perturbation clusters, we used well-defined phosphosites as pathway activation markers (Figure 5C) (Soste et al., 2014). Combining these pieces of information allows more detailed characterization of stress responses. For example, cluster 2 contains the high pH perturbations and shows unique activation dynamics for the Ipl1, Mck1 and Cla4/Ste20/Ypk2 kinases. Ipl1 kinase mediated phosphorylation of Dam1 S23 directly inhibits attachment of the kinetochore to the microtubules and Cla4/Ste20/Ypk2 mediated phosphorylation of Myo5 S357 is required for polarization of the actin cytoskeleton. Interestingly, pH changes have been associated with modulation of cell cycle (Orij et al., 2012) and actin polymerization (Wioland et al., 2019), which is likely induced by the stress response defined in cluster 2. We identified 57 phosphosites that showed selective regulation in one of the perturbation clusters (Figure 5D, Table S7). These phosphosites can serve as markers of the different stress responses and find application in investigating pathways and biological processes targeted and towards identifying stress responses in other phosphoproteomic datasets.

### Activity and architecture of the TOR signaling network

To investigate the organization of major signaling hubs we chose to focus on the TOR signaling network. We selected 9 kinases and phosphatases associated with TOR signaling (Dokládal et al., 2021) and matched these with sets of phosphosite targets curated from the literature to produce a list of 934 total phosphosites with which we investigated dynamics within the TOR network (Figure S5A, Table S8). Given the number of phosphosites available to analyze, we employed Weighted Gene Correlation Network Analysis (WGCNA) (Zhang and Horvath, 2005) for determining groups of co-regulated phosphosites and their dynamics (Langfelder and Horvath, 2008). The WGCNA analysis resulted in seven TOR subnetworks, which varied in size from 23 to 287 phosphosites (Figure 6A; Figure S5B-S5D, Table S8).

**Figure 6.**
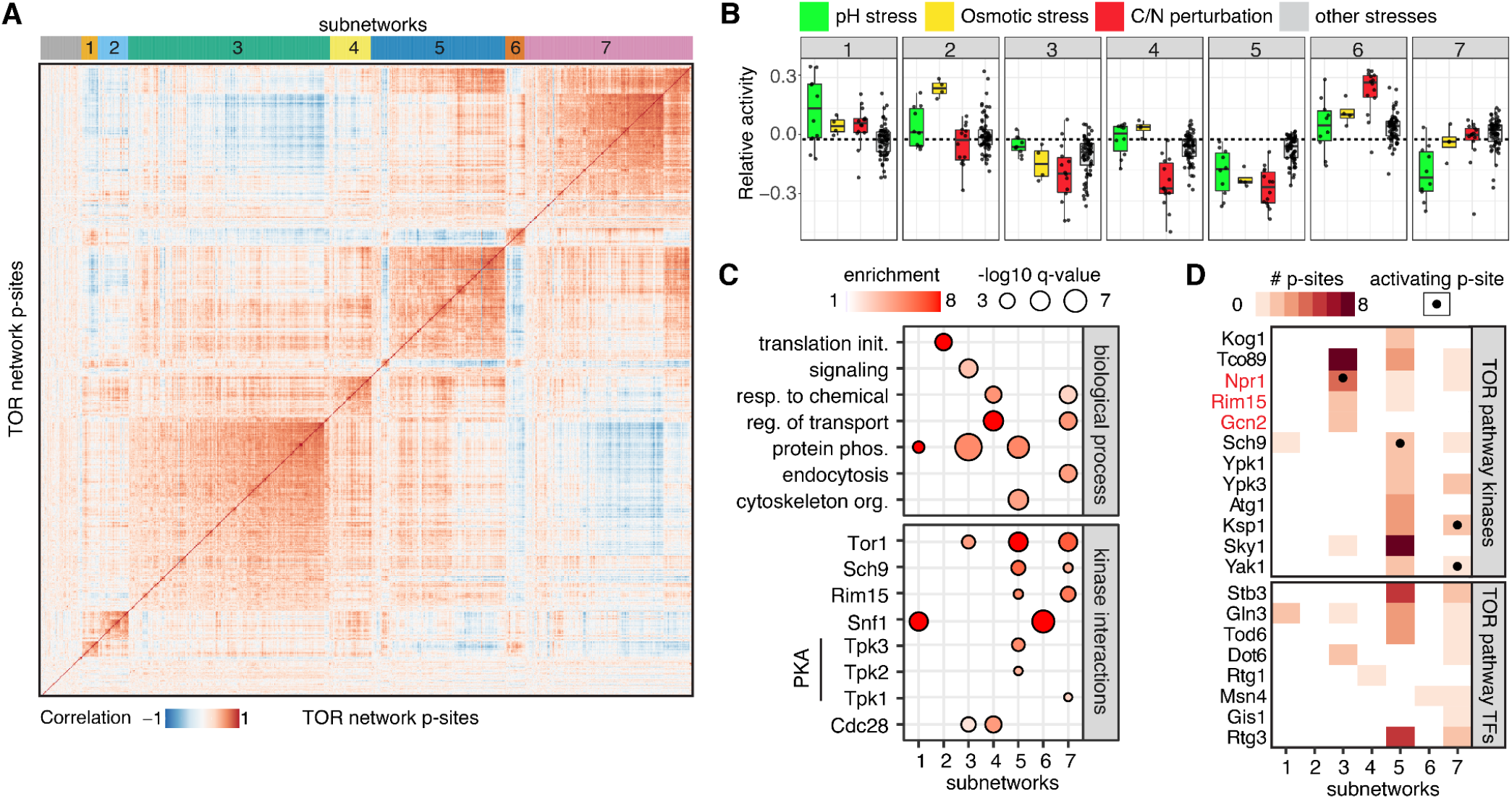
Regulatory dynamics of the TOR signaling network. **(A)** Correlation heatmap for targets of the TOR signaling network. Phosphosites (N=934) were clustered and broken into 7 subnetworks using WGCNA. **(B)** Activity of subnetworks across perturbations grouped by perturbation type. Subnetwork activity is defined by the first eigenvector of the singular value decomposition of each subnetwork’s expression across perturbations. **(C)** Significantly enriched (Fisher exact test q-value < 0.01) GO biological processes (top) and kinase-protein interactions (bottom) for phosphoproteins within each subnetwork. **(D)** Heatmap of phosphosite counts on TOR pathway kinases and transcription factors within each subnetwork. Kinases marked in red are regulated by the Tap42-PP2A/Sit4 phosphatase branch of the TOR pathway.

Since TOR pathway is a master regulator of cell growth and metabolic state, we tested how three major perturbation groups -C/N source changes, pH perturbation and osmotic stress– drive subnetwork dynamics by measuring subnetwork phosphorylation relative to untreated cells (Figure 6B). Strikingly, all subnetworks showed clearly different response patterns across tested perturbation classes. For example subnetwork 4 is exclusively responsive to nutrient limitations whereas subnetwork 7 is exclusively responsive to pH perturbations. We hypothesized that subnetworks represent different branches of the TOR pathway defined by differential activities of TOR associated kinases and phosphatases. To test this hypothesis, we integrated gene ontology, kinase-protein interaction, linear motif and phosphosite analyses (Figure 6C, Figure 6D and Figure S5E) with a literature curated pathway map (Figure S5F and Supplementary Text for detailed subnetwork analysis and interpretation). We found that subnetworks 3 and 5 showed close proximity to Tor1, with multiple lines of evidence suggesting that subnetwork 3 represents targets of the TORC1-Tap42-PP2A/Sit4 phosphatase branch and subnetwork 5 targets of the TORC1-Sch9 signaling branch. The two branches showed high sensitivity to most nutrient perturbations and osmotic stresses; however only the TORC1-Sch9 branch was sensitive to pH perturbations. Additionally, we were able to reveal subnetworks that likely resulted from pathway cross-talk. We found evidence for the exclusively pH-sensitive subnetwork 7 being at the downstream intersection of TORC1-Sch9 and PKA signaling pathways, and subnetwork 1 and 6 at the intersection of the SNF1 and TOR pathways. Interestingly, the analysis also identified co-regulated phosphosites on downstream effector protein targets. For example subnetwork 2 was strongly and exclusively activated by osmotic stress and contained multiple translation initiation factors, suggesting that these phosphosites might be part of the osmotic stress-induced translation inhibitory response.

It has not been possible so far to systematically assess how different stimuli affect a single pathway and leverage this information to derive pathway organization at the phosphosite level. Our analysis of the TOR pathway suggests that, while defining kinase-substrate relationships is important, measuring how phosphosites are co-regulated across perturbations allows understanding signaling network wiring. Our phosphoproteomic resource and the analysis approaches outlined here can be applied to any known or newly defined pathway to determine functional pathway organization into subnetworks and perturbation-specific activation states.

### Functional organization of the stress responsive phosphoproteome

Next, we tested if functional phosphoproteome organization can be inferred from co-regulated phosphosites alone, i.e. without seeding kinases-substrate relationships. We employed phosphosite quantification of the core phosphoproteome and performed analysis in low-dimensional Uniform Manifold Approximation and Projection (UMAP) space. UMAP embedding of individual phosphoproteomes showed clear separation of broader perturbation types (Figure 7A) and matched underlying pathway activities (TOR, HOG, SNF1), as judged by pathway activation markers across embedded phosphoproteomes (Figure 7B). Using autocorrelation statistics, we defined 2,191 phosphosites that significantly vary across samples in UMAP space. We performed UMAP embedding of the quantitative values of these phosphosites across perturbations, and conducted community analysis to identify groups of co-regulated phosphosites. This resulted in 29 major phosphosite modules that contained between 20-140 phosphosites each (Figure 7C, Figure S6A and S6B). Strikingly, phosphosite motif class was a main factor for separation in UMAP space (Figure 7D, Figure S6C), indicating that co-regulated phosphosites are modified by the same or closely-related kinases. We characterized co-regulated phosphosites within each module by performing a series of analyses, including enrichment for motif, phenotype, gene ontology, kinase-protein interactions, and kinase-substrate associations, as well as integrating modules as STRING networks. Additionally, we established relationships among the different modules by comparing: (1) overall abundance of phosphosites within a module; (2) aggregated abundance of phosphosites across perturbations; and (3) significantly regulated phosphosites versus untreated within a module. Characterization of phosphosites within each module was assembled in a comprehensive catalog (Table S9). This integrated resource can greatly help identify new potential pathway substrates, as well as greatly assist other researchers with pre-selecting pathways and conditions that might regulate a specific phosphosite of interest for further investigation. To showcase the kind of knowledge that can be generated from the data, we chose a few distantly and closely related modules for more detailed analysis (Figure 7E and 7F, Table S9 and Supplementary Text for detailed analysis of selected modules).

**Figure 7.**
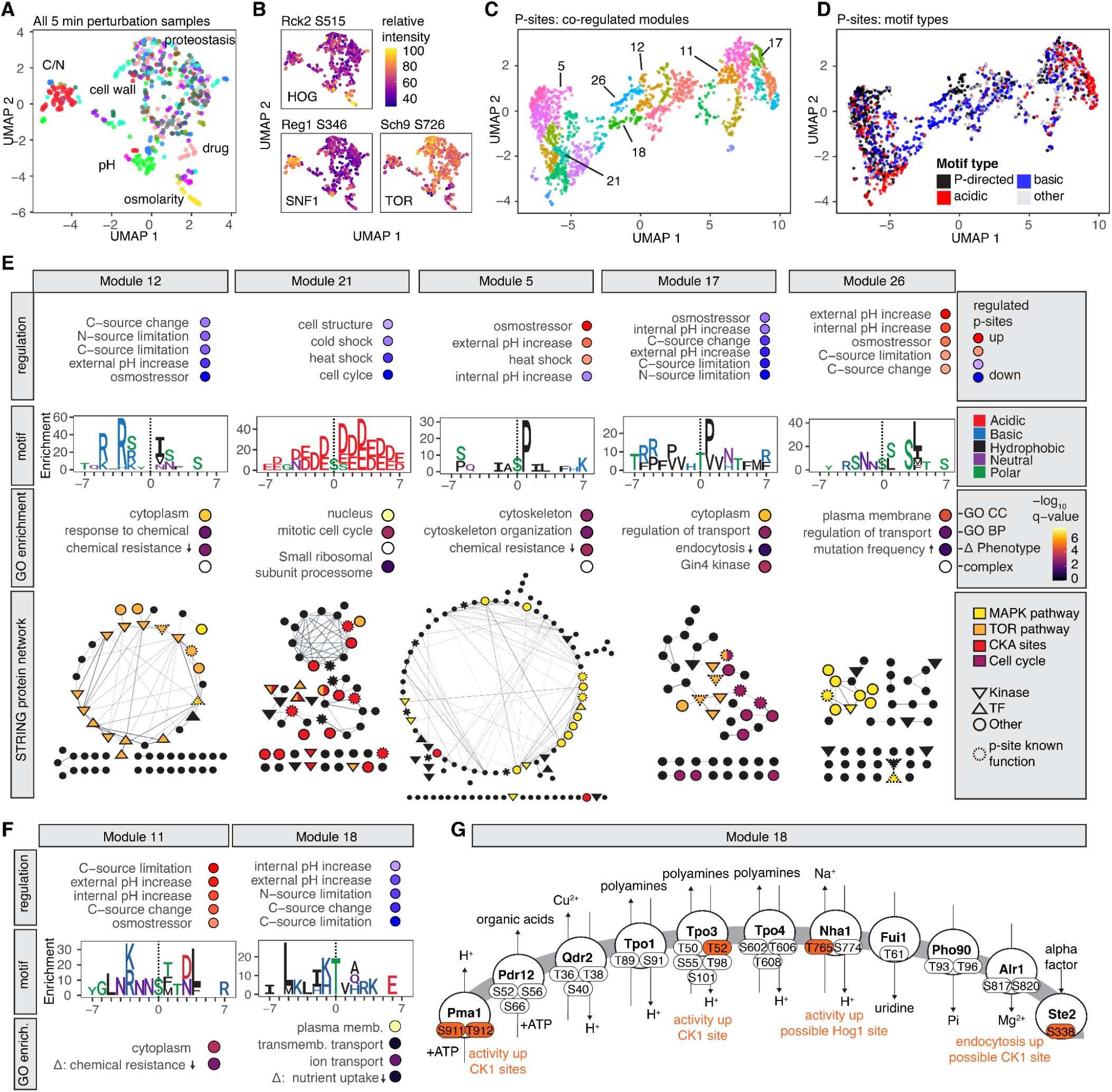
Functional organization of the stress responsive phosphoproteome revealed through dimensionality reduction and co-regulation analysis. **(A)** UMAP embedding of individual samples based on their underlying phosphoproteome. Samples are color coded based on perturbation type. The most common perturbation types within different areas of the UMAP are indicated. Perturbations are color coded according to Figure 1. **(B)** Relative intensity of pathway activity marker phosphosites overlaid on UMAP from (A). **(C)** UMAP embedding of 2,191 phosphosites based on their quantitative profiles across samples. Modules of co-regulated phosphosites were defined using Louvain community analysis. Modules are color-coded and annotated if they are discussed in the text. **(D)** Same UMAP as in (C) color-coded by motif type of phosphosites. **(E)** Functional characteristics of co-regulated phosphosites within five selected modules. Top row: perturbation types with most up- or down-regulated phosphosites within a module. Middle rows: motif enrichment for phosphosites within a module versus all phosphosites and GO/phenotype enrichment. Bottom row: STRING network of proteins within the module. **(F)** Regulation, phosphosite motif, and GO/phenotype enrichment for modules 11 and 18. **(G)** Plasma membrane phosphoproteins from module 18 annotated with functions, co-regulated phosphosites and phosphosites with known function or regulation (orange).

First, co-regulation analysis allowed us to recover known kinase-substrate relationships and identify novel kinase substrates. For example, we identified module 12 to be downstream of TORC1-Sch9, characterized by a basic phosphorylation motif, a high enrichment for Tor1 and Sch9 kinase-protein interactors, and high sensitivity to nutrient perturbations. Indeed, 30% of these phosphosites were annotated to the TORC1-Sch9 signaling branch in our previous TOR signaling network analysis (TOR subnetwork 5). Importantly, 55% of phosphosites in module 12 did not contain any previous connection to the TOR pathway but are likely newly identified TORC1-Sch9 targets. Similarly module 21 had multiple lines of evidence for being targets of casein kinase 2 and Module 11 for Snf1 (Figure 7E).

Second, we were able to resolve differentially regulated phosphosites with similar consensus motifs. Module 5 and module 17 showed both a proline-directed motif, however module 5 is upregulated in osmotic stress and heat shock whereas module 17 shows the opposite regulation and is additionally sensitive to nutrient perturbations. We could associate module 5 to MAPK cascades and module 17 to down-regulated activity of the master cell cycle kinase Ccd28 based on the protein-protein interaction network (Figure 7E).

Third, we identified regulatory functions and potential novel targets for understudied kinases. Phosphosites of module 26 showed evidence for action of Kin1/Kin2 kinases (orthologous to MARK/PAR-1, AMPK family members) based on phosphorylation motif and kinase-phosphoprotein interactions. Additionally phosphoproteins in this module were enriched for plasma membrane proteins and regulation of transport (Figure 7E). Kin1/2 kinases have been reported to localize to the plasma membrane, be associated with HOG signaling and activated upon cellular stress, and regulate exocytosis (Ghosh et al., 2018; Romanov et al., 2017). Taken together, this indicates a function for Kin1/2 in inducing plasma membrane protein phosphorylation during cell wall stress.

Fourth, we revealed effector modules consisting of co-regulated phosphosites on proteins with similar functions. For phosphoproteins in module 18 we found a striking functional enrichment of plasma membrane proteins that are involved in ion homeostasis, transport and required for proper nutrient uptake (Figure 7F and 7G). Several identified phosphosites on these proteins are known to increase their transport activity or are required to stabilize the protein at the plasma membrane and had evidence for activity of plasma membrane bound casein kinase I. Taken together, the phosphosites contained in this module likely orchestrate ion homeostasis upon intracellular pH changes through a common mechanism.

In summary, we demonstrate that co-regulation analysis of phosphosites across many perturbations can be used to reveal functional organization of the phosphoproteome. In addition to the quantitative phosphoproteomic atlas, we created an information-rich catalog of modules that contain co-regulated phosphosites with functional and/or regulatory connections (Table S9) that can serve as a starting point to assemble novel signaling pathways and interrogate perturbation-specific protein functions. We anticipate that the provided resource will facilitate hypothesis generation for mechanistic protein-phosphosite and signaling studies in the future.

## Discussion

A global view of how signaling networks link environmental fluctuations to cellular processes is currently lacking. High-dimensional perturbation-signaling maps will be key to unbiasedly understand the complex network interactions that govern cell function. In this study, we demonstrated that phosphoproteomic measurements of a wide variety of cellular responses to perturbations can reveal activation states and make considerable contributions to understanding the functional organization of signaling networks.

We provide the deepest resource of yeast phosphosites with quantifications under standard growth conditions and upon 5 min exposure to different perturbations. A considerable number of phosphosites (~40%) within our reference phosphoproteome have not been reported before. We found that phosphosites occur more frequently on protein termini and disordered protein regions; and we identified ~300 proteins with >30 phosphosites, suggesting a potential role in regulating liquid-liquid phase separation through bulk electrostatic changes (Yamazaki et al., 2022).

Regulated phosphosites were more likely to impact protein function and had a higher probability of occurring at conserved regions within proteins. Analyzing yeast paralogs showed that a high number of phosphosites in one paralog is substituted by a phosphomimetic residue in the other paralog. Such an evolutionary switch might lead to functional diversification by introducing phosphorylation-dependent and condition-specific protein functions. Future evolutionary analysis of phosphosites can benefit from including conditional regulation along with sequence conservation (Studer et al., 2016).

Regulation of the phosphoproteome agreed well with prior results of phosphorylation for individual stresses or targeted pathways (Figure 2). For example our resource recapitulates regulation of many canonical TOR pathway targets in nutrient limited conditions (Figure 3). However, quantifying responses of the same signal transduction network across many different perturbations gave us the power to dissect layers of regulatory connections between signaling events that remained hidden in low-dimensional studies. We were able to identify TOR subnetworks by grouping phosphosites of the TOR network according to their regulatory profile and connecting them to the activity of specific TOR kinases and phosphatases. These TOR subnetworks showed distinct responses to perturbations (Figure 6). Furthermore, dimensionality reduction and co-regulation analysis identified many phosphosites and proteins that have not been previously associated with TOR signaling and positioned them within the different subnetworks (Figure 7). Our analysis of the well-studied TOR pathway revealed a much broader involvement of TOR in the signaling response to stress than anticipated as well as the multiple ways in which the TOR network intercalates with other stress-responsive pathways.

Systematic kinase deletion studies (Bodenmiller et al., 2010; Li et al., 2019) have shown widespread changes to the proteome and phosphoproteome owing to the high interconnectivity of the signaling network. We find that even for highly selective kinases, target phosphosites partition into distinctly regulated subsets. Thus high-dimensional phosphoproteomic studies measuring early response to perturbation can enhance our understanding of direct kinase action, revealing at a systems-level, previously unappreciated subnetworks, subnetwork intersections, and subnetwork sensitivities.

Our data show that dephosphorylation is a phosphoproteomic response common to many types of perturbations when compared to untreated, exponentially growing cells. Exponential growth requires the consistent and combined activity of several major nutrient responsive and cell cycle promoting pathways (TOR, PKA, CDK). These pathways are induced if environmental conditions are favorable, and they control the activities of large sets of kinases and phosphatases to promote biosynthetic processes and repress catabolic processes. We identify hundreds of down-regulated phosphosites that have been connected to these pathways underscoring the role of phosphatases in stress response. Whereas nutrient limitations lead to a consistent and strong dephosphorylation of the TOR and PKA signaling networks, many other perturbations also lead to their dephosphorylation. Through co-regulation analysis, we find points of pathway cross-talk that might amplify this early stress response. Examples of cross-talk include the regulation of proline-directed phosphosites on TOR effector kinases that mirror activities of MAPK pathways; a signaling axis that is exclusively responsive to pH perturbation at the intersection of the TOR and PKA pathway; and reciprocal phosphorylation by TOR and SNF1 on shared substrates.

Our integrated analysis approaches uncovered many unknown or poorly-characterized stress response trajectories, such as a group of plasma membrane transporters that are presumably deactivated upon pH and nutrient perturbations; Hal5 kinase as an integrator of a multitude of stress responses; a role for Kin1/2 kinases in the pH stress response; and deactivated casein kinase 2 in heat and cold shock. As showcased throughout this study, the created resource will be invaluable for stimulating novel hypotheses: (1) for phosphosite and protein centric analyses the resource can be mined for phosphosites that occur on specific proteins, how these phosphosites are regulated and by what potential kinase or pathway; (2) for pathway centric analyses known or newly identified signaling modules are provided and can be interrogated for perturbation specific activation states; (3) for perturbation centric analyses, shared and specialized phosphorylation response programs were defined and can be used to explore regulated cellular processes.

In order to advance analysis of our data set and to assist scientists in generating hypotheses, we provide detailed analysis on identified signaling modules (Table S9) and have developed a web-based analysis tool (https://yeastphosphoatlas.gs.washington.edu) that allows easy access of regulatory profiles of individual or groups of phosphosites across perturbations. Interest in understanding phosphorylation signaling networks across many perturbations or large sample cohorts in basic and translational research is rapidly growing and the experimental and computational phosphoproteomic approaches developed as part of this study can be used to guide and expand these efforts.

## Methods

### Cellular treatments and harvest

The S288C-derivative BY4741 (MATa his3Δ1 leu2Δ0 met15Δ0 ura3Δ0) *Saccharomyces cerevisiae* strain was used for all treatments, except for the diploid condition where BY4743 was used. Unless mentioned otherwise, all treatments were done as described next. Single colonies were picked from a fresh plate and grown overnight, shaking at 30°C in Synthetic Complete media with 2% Glucose (SC+C). Next day, 50ml cultures were inoculated in SC+C at an OD600 of 0.1 and grown for ~5h until they reached an OD600 of 0.6. Treatments were performed on these cultures. For treatments that required a media exchange, yeast were filter-purified and filters were incubated in prewarmed new media. Each culture was treated as an independent replicate and treatments were performed shaking at 30°C for exactly 5 min (or as described for specific treatments). A description of exact treatment conditions can be found in Table S1. To stop treatments, cells were metabolically arrested and harvested by adding 100% (w/v) trichloroacetic acid (TCA) directly to the liquid culture to a final concentration of 10% TCA. Cultures were incubated on ice for 10 min, centrifuged, decanted, washed once with 100% acetone at −20°C, centrifuged, decanted, washed with ice-cold water, centrifuged and decanted. Cell pellets were snap-frozen in liquid nitrogen, and stored at −80°C until lysis.

### Cell lysis, protein reduction, and alkylation

Frozen cell pellets were resuspended in a lysis buffer composed of 8 M urea, 75 mM NaCl, and 50 mM HEPES pH 8. Cells were lysed by 4 cycles of bead beating (30-s beating, 1-min rest on ice) with zirconia/silica beads followed by clarification by centrifugation. Protein concentration was measured for every lysate by BCA assay and adjusted to 1 mg per ml. Proteins were reduced with 5 mM dithiothreitol (DTT) for 30 min at 55°C and alkylated with 15 mM iodoacetamide for 15 min at room temperature in the dark. The alkylation reaction was quenched by incubation with additional 10 mM DTT for 15 min at room temperature. Lysates were stored at −80°C until further processing.

### Proteomic and phosphoproteomic sample preparation based on R2-P2

Lysates were scrambled across 96-deep well plates and biological replicates were blocked from being on the same 96-well plate (Figure S1A). Each plate contained 4 samples consisting of the same pooled lysate to assess sample preparation reproducibility between 96-well plates. To purify proteins and perform digestions, 96-well plates were processed using the R2-P1 (Rapid-Robotic proteomics) protocol implemented on a KingFisher™ Flex (Thermo Fisher) magnetic particle processing robot as established by our group before (Leutert et al., 2019). Briefly, for every sample, 250 μl lysate (250 μg protein) was combined with 250 μg carboxylated paramagnetic beads (1:1 mix of hydrophiclic and hydrophobic Sera-Mag SpeedBead Carboxylate-Modified, GE Life Sciences) and 750 μl ethanol. Protein-bead binding occurred for 30 min and protein-bead complexes were purified by moving through 4 wells containing 500 μl 80% ethanol. Proteins were eluted and digested in 500 μl 25 mM ammonium-bicarbonate buffer containing 2.5 μg trypsin (sequencing grade, Promega) for 5 h at 30°C. Digests were acidified directly in 96-well plates, 5% of the sample was taken out for total proteomic analysis and all samples were lyophilized in a speedvac overnight. The next day, samples were resuspended within the 96-well plate in 80% ACN (acetonitrile), 0.1% TFA (trifluoroacetic acid), clarified by centrifugation and R2-P2 (Rapid-Robotic Phosphoproteomics) was performed on a Kingfisher Flex (Leutert et al., 2019). Briefly, phosphopeptides were enriched with Fe^3+^-NTA magnetic beads, (PureCube Fe-NTA MagBeads, Cube Biotech), washed three times with 80% ACN, 0.1% TFA, eluted in 50% ACN, 2.5% NH_4_OH, acidified, lyophilized in a speedvac and store at −20°C until MS-analysis. Sample preparation metadata including order and batches is available in Table S1.

### Ultra-deep reference phosphoproteome sample preparation

For the ultra-deep reference phosphoproteome, a pooled sample (only 5-min perturbations) was created by combining equal amounts of lysates from each sample. In-solution digestions were performed on lysates containing 5 mg of protein using either Trypsin (Promega), Chymotrypsin (Promega), Glu-C (Promega) or Lys-C (Wako chemicals) according to manufacturer protocols. Peptides were desalted on C18 SepPak cartridges. Phosphopeptides were enriched using R2-P2.

Offline pentafluorophenyl (PFP) reverse-phase chromatography was performed on phosphopeptides derived from the different digests individually using a XSelect HSS PFP 200 × 3.0 mm; 3.5 μm column (Waters) as described (Grassetti et al., 2017). 48 fractions were collected and combined into 12 pooled fractions and lyophilized.

Strong cation exchange (SCX) chromatography was performed on tryptic digest using a polysulfoethyl A, 200 × 4.6mm; 5μm, 300A column (PolyLC) and two buffers: (A) 10 mM ammonium formate, 0.05% formic acid, 25% ACN and (B) 500 mM ammonium formate, 0.05% formic acid in 25% ACN. Peptides were fractionated with a gradient ranging from 5% buffer B to 100% buffer B, 12 fractions were collected, lyophilized and phosphopeptides enriched with R2-P2 (Villén and Gygi, 2008).

### Mass spectrometry data acquisition

Lyophilized peptide and phosphopeptide samples were dissolved in 4% formic acid, 3% acetonitrile and analyzed by nLC-MS/MS. Peptides were loaded onto a 100 μm ID × 3 cm precolumn packed with Reprosil C18 3 μm beads (Dr. Maisch GmbH), and separated by reverse-phase chromatography on a 100 μm ID × 35 cm analytical column packed with Reprosil C18 1.9 μm beads (Dr. Maisch GmbH) and housed into a column heater set at 50°C.

DIA-MS measurements for phosphoproteomic and total proteomic samples were performed on an Orbitrap Exploris 480 Mass Spectrometer (Thermo Fisher) equipped with an Easy1200 nanoLC system (Thermo Fisher). Peptides were separated by a 60-min effective gradient ranging from 6 to 30% acetonitrile in 0.125% formic acid. DIA measurements were set-up and optimized according to (Pino et al., 2020) and the DIA window scheme is visualized in Figure S1A. For DIA measurements, we acquired 30 × 24-m/z (covering 438-1170 m/z for phospho samples and 363 to 1095 m/z for total proteome samples) precursor isolation window MS/MS DIA spectra (30,000 resolution, AGC target 1e6, auto inject time, 27 NCE) using a staggered window pattern and optimized window placements. Precursor spectra (60,000 resolution, standard AGC target, auto inject time) were interspersed every 30 MS/MS spectra. For total proteome measurements we acquired 30 × 24-m/z (covering 363-1095 m/z) precursor isolation window MS/MS DIA spectra with otherwise similar settings as the phosphoproteome measurement.

DDA-MS measurements for the deep phosphoproteome and spectral libraries were either acquired on an Orbitrap Exploris 480 mass spectrometer (Thermo Fisher) or on an Orbitrap Eclipse Tribrid mass spectrometer (Thermo Fisher) equipped with an Easy1200 nanoLC system (Thermo Fisher) using 60-min effective gradients and performing all measurements within the Orbitrap.

### Mass spectrometry data analysis

The *S. cerevisiae* S288C reference protein fasta database containing the translations of all 6713 systematically named ORFs, except “Dubious” ORFs and pseudogenes created on 05/11/2015 by SGD (https://www.yeastgenome.org/) was used for all searches.

DDA data was searched with Comet (2019.01.2) (Eng et al., 2013). The precursor mass tolerance was set to 20 ppm. Constant modification of cysteine carbamidomethylation (57.021463 Da) and variable modification of methionine oxidation (15.994914 Da) were used for all searches, and additional variable modification of serine, threonine, and tyrosine phosphorylation (79.966331 Da) was used for phosphopeptide samples. Search results were filtered to a 1% FDR at PSM level using Percolator (Käll et al., 2007). Phosphosites were localized using an in-house implementation of the Ascore algorithm (Beausoleil et al., 2006). Phosphosites with an Ascore > 13 (p-value < 0.05) were considered confidently localized.

For spectral library generation and spectral library searches of DIA data, Spectronaut v.15 (Biognosys) was used. A hybrid phosphopeptide spectral library was generated by searching all DDA data encompassing the deep yeast phosphoproteome coming from tryptic digests together with DIA data encompassing the quantitative phosphoproteomic measurements. Standard search parameters were used, including fixed modification of cysteine carbamidomethylation and variable modification of methionine oxidation and serine, threonine, and tyrosine phosphorylation. A PSM and peptide FDR cutoff of < 0.01 and a PTM localization site confidence score cutoff of > 0.75 were chosen.

For the spectral library searches of the phosphoproteomic DIA data standard spectral library search setting with following adjustment were chosen: decoy limit strategy was set to dynamic with a library size fraction of 0.1, but not less than 5000, a precursor FDR cutoff of < 0.01 was enforced by choosing the data filtering setting “Qvalue”, no imputation or cross run normalization was performed, a PTM localization site confidence score cutoff of > 0.75 was chosen, multiplicity was set to false, and PTM consolidation was done by summing. Raw files were searched in batches of 100 files and combined all together using the “SNE combine workflow” in Spectronaut to merge the identification results of individual batches in a FDR controlled manner. Spectral library searches of the total proteomic DIA data were done in similar manner as the phosphoproteomic searches, with a total proteome spectral library.

Sample outliers due to obvious LC, MS, or sample preparation problems as judged by LC, MS, or sample quality control parameters (Figure S1C) were excluded from analysis.

### Differential expression analysis

Quantitative phosphoproteomics analysis was performed at the phosphosite level across the data by summing peptide quantification values and median normalization across individual samples. As described in the text, a stringent filter for phosphosites which were present in at least 50% of all samples was applied and remaining missing values were imputed using the QRILC algorithm from the imputelcmd R package (https://cran.rstudio.com/web/packages/imputeLCMD/index.html). For differential expression analysis, surrogate variable (SV) analysis was performed on all samples using the SVA R package (https://bioconductor.org/packages/release/bioc/html/sva.html), with the number of significant SVs determined by the package. Differential expression was determined using LIMMA (https://bioconductor.org/packages/limma/) on all samples at once with a model matrix including both the treatment for each sample and the individual SVs to correct batch information. Significant differential expression was then calculated for each treatment against the untreated samples, and p-values were corrected globally using Benjamini-Hochberg correction. Finally, a phosphosite was called as significantly regulated in a treatment if its absolute log2 fold change against untreated samples was at least 1 and its adjusted p-value was less than 0.05.

### Co-regulation analysis and UMAP dimensionality reduction

For phosphosite-by-phosphosite correlation and UMAP analysis of the data, a second dataset was created from the filtered data with R2-P2 batch effects directly corrected using the COMBAT method from the SVA package (Table S5).

Weighted gene correlation network analysis (WGCNA) was performed using the WGCNA package (Zhang and Horvath 2005). The first step of WGNCA is to pick an integer power to raise the Pearson correlation matrix to, which is then used to transform the matrix before calculation of the topological overlap matrix. This soft threshold was chosen as the first power which resulted in a scale free model R^2^ passing 0.8, as recommended by the method’s authors.

To project individual sample phoshoproteomes and phosphosite quantifications into two dimensions, we performed dimensionality reduction with the UMAP algorithm (McInnes et al., 2018) using functions in the Monocle3 R package (Cao et al., 2019; Qiu et al., 2017; Trapnell et al., 2014) and approaches outlined in (Dorrity et al., 2020). Core Phosphosite quantifications in individual samples were given as input to Principal Component Analysis (PCA) using the Monocle 3 wrapper function “preprocess_cds”, remaining sample preparation batch effects were subtracted using the “align_cds” function, and the top 50 principal components were then used as an input to UMAP using the “reduce_dimension” wrapper function with default parameters (except: umap.min_dist = 0.1, umap.n_neighbors = 10L) for generating 2D projections of the data. Phosphosites that vary across samples in UMAP space were identified using spatial autocorrelation statistics (Moran’s I) as implemented in the function “graph_test” with default parameters (except: k=10) and filtered for a q-value < 0.05 and morans_I > 0.2 (effect size). To group phosphosites that vary in a similar way across perturbation into modules, the “find_gene_modules” function was applied using default parameters (except: k = 15, umap.n_neighbors = 10L) to run UMAP on phosphosites and grouped them into co-regulated modules using Louvain community analysis.

### Other bioinformatic analyses

If not specified otherwise, R version 4.1.0 (https://www.r-project.org/) with the “tidyverse” package collection (https://www.tidyverse.org) was used for all analyses. Heatmaps were done using the “pheatmap” R package (https://cran.r-project.org/web/packages/pheatmap/index.html). For motif sequence logo plots enrichment was calculated using the R package “dagLogo” (https://www.bioconductor.org/packages/release/bioc/html/dagLogo.html) and motifs were plotted using the R package “ggseqlogo” (https://cran.r-project.org/web/packages/ggseqlogo/index.html)

Gene ontology enrichment analysis was performed by annotating phosphopeptides with the corresponding protein annotation terms and using the whole yeast proteome (protein level) or the deep phosphoproteome (phosphosite level) as background if not otherwise stated. For enrichment analysis Fisher exact test with Benjamini–Hochberg multiple-hypothesis correction was applied and filtered for q-values < 0.01. The categories used for the annotation of proteins were GO Biological Processes, GO Cellular Components, GO Molecular Functions, protein complexes and phenotypes of gene deletion mutants, all downloaded from SGD (https://www.yeastgenome.org/ on 04/05/2022. Kinases and phosphatase target annotation were downloaded from BioGrid (https://thebiogrid.org on 10/23/2020) and completed with data from (Dokládal et al., 2021; Goldman et al., 2014; Hu et al., 2019; Li et al., 2019; Rubenstein and Schmidt, 2007; Soste et al., 2014; Stark et al., 2010; Velázquez et al., 2020; Wagih et al., 2018).

Cytoscape 3.9.1 was used to create and plot STRING networks of phosphoproteins at a minimum required STRING interaction score of 0.4.

For all boxplots, the lower and the upper hinges of the boxes correspond to the 25% and 75% percentile, and the bar in the box to the median. The upper and lower whiskers extend from the largest and lowest values, respectively, but no further than 1.5 times the IQR from the hinge.

## Supplementary material

Supplementary Information: Supplementary Figures and Supplementary Text.

Table S1: Treatment, sample preparation and measurement information for all analyzed samples, related to all Figures.

Table S2: Ultra-deep reference yeast phosphoproteome, related to Figure 1 and 2.

Table S3: Complete and uncorrected, quantitative phosphosite datasets, related to Figure 1.

Table S4: Proteome quantitation of 30 selected perturbations and differential expression of proteins versus untreated. Related to Figure 3.

Table S5: Corrected quantitative core phosphoproteome dataset, related to Figure 1 and 7.

Table S6: Differential expression of phosphosites in all perturbations versus the untreated control. Related to Figure 3 and 4.

Table S7: Perturbation-specific phosphosites. Related to Figure 5.

Table S8: Phosphosites associated with TOR subnetworks. Related to Figure 6.

Table S9: Catalog of characterized, co-regulated phosphosites, assembled in modules. Related to Figure 7.

## Data availability

The mass spectrometry proteomics data have been deposited to the ProteomeXchange Consortium via the PRIDE (Perez-Riverol et al., 2022) partner repository with the dataset identifiers PXD035029 for the ultra-deep reference phosphoproteomic DDA data, PXD035050 for the quantitative phosphoproteomic DIA data, and PXD034997 for the quantitative proteomics DIA data.

## Code availability

The code for processing and analyzing the phosphoproteomic data is available at: https://github.com/Villen-Lab/YeastPhosphoAtlasAnalysis. The code for the web resource is available at: https://github.com/Villen-Lab/YeastPhosphoAtlas.

## Acknowledgments

We thank Matthew Berg, Kyle Hess, Alexander Hogrebe, Julian Ramos, Ian Smith, Maitreya Dunham and members of the Villén Lab for useful discussions and feedback. We thank Life Science Editors for editing services (www.lifescienceeditors.com). These studies were supported by the National Institutes of Health grants R35GM119536 and R01AG056359. M.L. was supported by the Swiss National Science Foundation grants P2ZHP3_181503 and P400PB_194379. A.S.B. was supported by the NIH training grant T32LM012419.

## Author contributions

M.L. and J.V. conceived the project. M.L conducted experiments with assistance from N.K.F and R.A.R.-M. M.L. and A.S.B. analyzed the data. A.S.B. created the website. M.L. and J.V. wrote the manuscript and all authors edited it.

