## Supplementary Information for "The regulatory landscape of the yeast phosphoproteome"

#### Index

| | |
|---|---|
| Page 2-12 | Supplementary Figures S1 - Figures S6 |
| Page 13-17 | Supplementary Text |
| Page 18-19 | Supplementary References |

#### Supplementary Figures

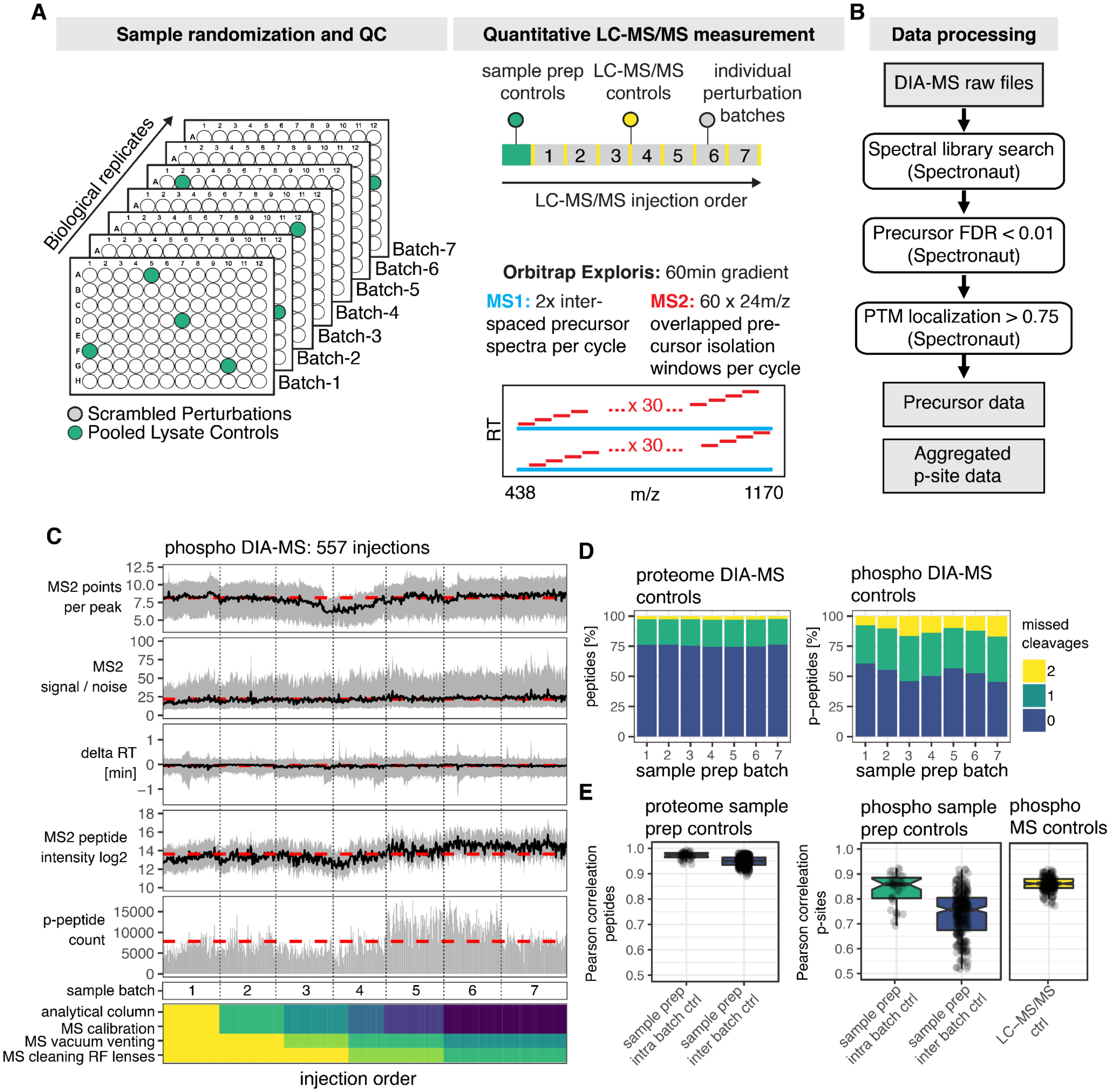


**Figure S1 – Experimental setup and quality control of sample preparation and mass spectrometry measurements, related to Figure 1**

**(A)** Sample randomization and quality control: Cell lysates were scrambled across 96-well plates and biological replicates were assigned to different 96-well plates. On each plate, 4 samples containing the same pooled lysate were included to assess sample preparation reproducibility between 96-well plate batches. Quantitative LC-MS/MS measurement: Proteomic and phosphoproteomic sample preparation controls in each batch were assessed first and then individual sample batches were measured. Performance of LC-MS/MS was regularly assessed between and within batches using a pooled phosphopeptide sample. DIA-MS was performed on an Orbitrap Exploris mass spectrometer using a method with a 60-min effective gradient and staggered wide-window DIA as depicted. **(B)** Overview of data processing workflow for DIA files that includes spectral library searches, application of a global precursor FDR (<0.01) and a PTM localization filter (>0.75) using Spectronaut. Quantifications were aggregated to the phosphopeptide and phosphosite level. **(C)** Overview of different quality control parameters tracked across all DIA-MS runs for phosphoproteomic sample injections in chronological order. The dashed red lines indicate median values across all injections. The black line indicates mean values within an injection and the gray area depicts the area between the 25th and 75th percentile. Numbers of phosphopeptide identification are shown as a bar plot. The two bottom panels show sample preparation batches and different colors denote different cycles of LC-MS/MS maintenance. **(D)** Missed cleavage rates of identified peptides and phosphopeptides across the different sample preparation batches. **(E)** Boxplots of Pearson's correlation coefficients from pairwise comparisons of individual injections. Pooled lysate controls processed within or between different 96-well plates for the proteome (left) and phosphoproteome (middle) and phosphoproteomic control measurements of the same pooled sample across the whole experiment (right) are shown.

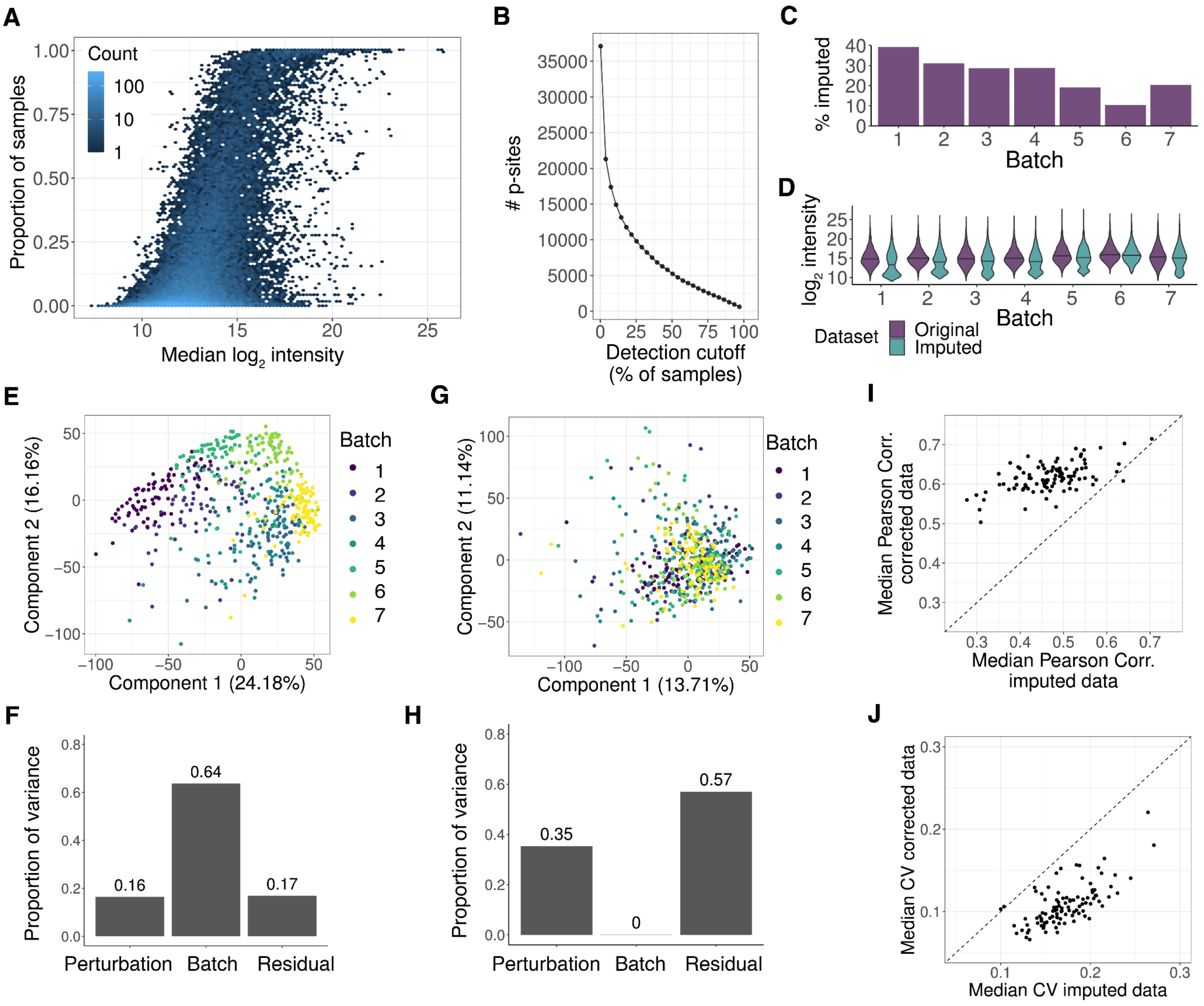


**Figure S2 – Missing data imputation and batch correction of the core phosphoproteome, related to Figure 1**

**(A)** Proportion of all samples where a phosphosite was detected vs the median $\log_2$ intensity of the phosphosite in the remaining samples before imputation and batch correction. Color indicates the number of phosphosites falling within a bin. The observed weak logistic trend implies that low peptide abundance is in part at play for missing quantifications. **(B)** Number of phosphosites remaining in the dataset after application of increasingly stringent cutoffs on the percent of samples in which phosphosites are detected. **(C)** Percentage of imputed phosphosites across batches after filtering out phosphosites which were not present in at least 50% of all samples. **(D)** Distribution of phosphosite intensities before and after imputation for each sample batch, with the median intensity for the batch displayed as a horizontal line. **(E)** PCA of phosphosite quantifications per sample colored by sample batch after phosphosites were filtered for missingness and imputed. **(F)** Principal Variance Component Analysis (PVCA) on the same data as E) demonstrating the proportion of variance explainable by the perturbation, the sample batch, and a residual component. **(G)** PCA of phosphosite quantifications per sample colored by sample batch after phosphosites were corrected using ComBat. **(H)** PVCA on the same data as (G) demonstrating the proportion of variance explainable by the perturbation, the sample batch, and a residual component. **(I)** Scatter plot of median Pearson correlation between samples from the same perturbation across batches before (imputed data) versus after batch correction (corrected data). **(J)** Scatter plot of the median coefficient of variation across phosphosites for each perturbation before (imputed data) versus after batch correction (corrected data).

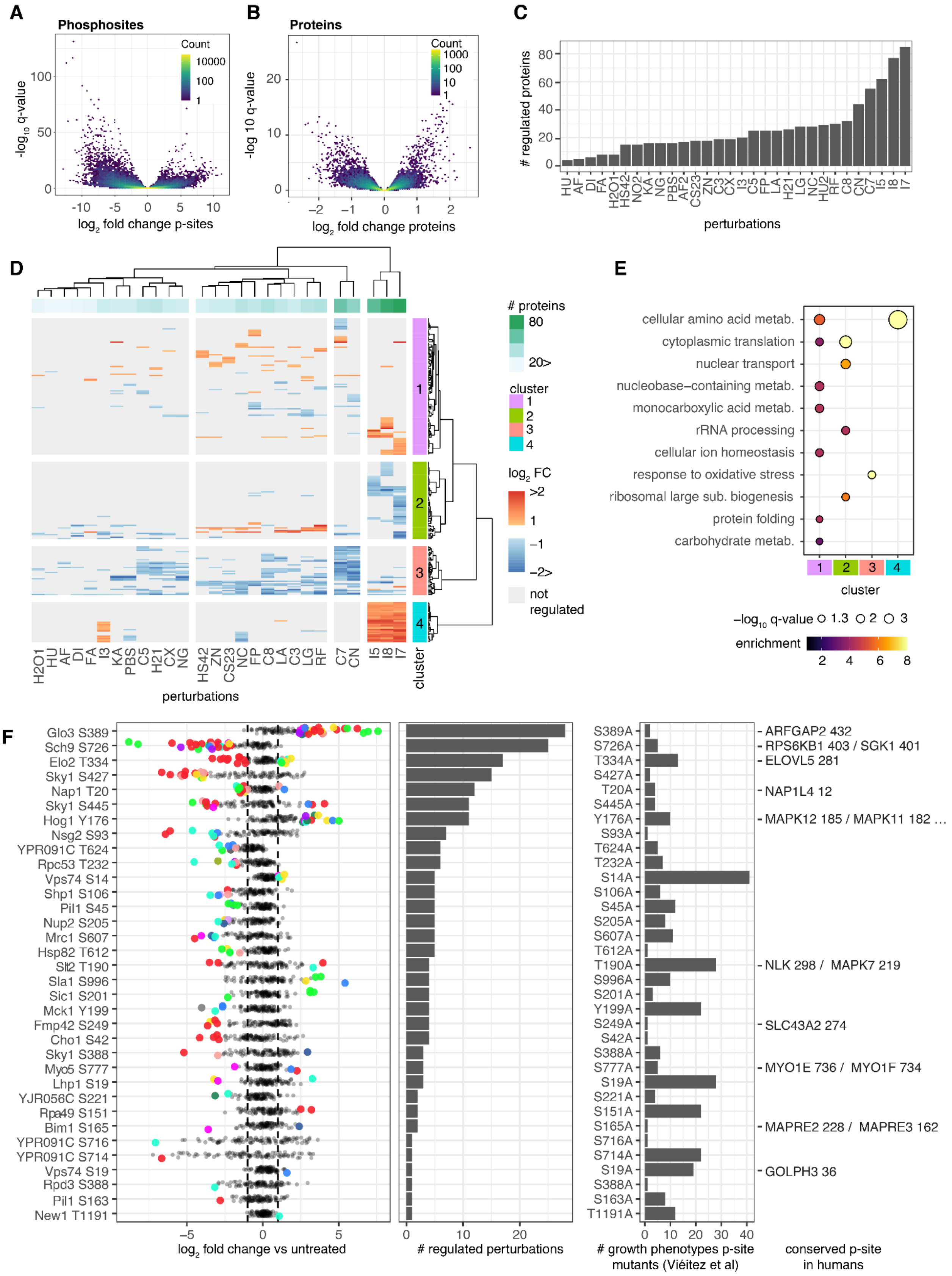
A
Phosphosites
Count
10000
100
1
−log10 q-value
log2 fold change p-sites
B
Proteins
Count
1000
100
10
1
-log 10 q-value
log2 fold change proteins
C
### regulated proteins
HU AF DI FA H2O1 HS42 NO2 KA NG PBS AF2 CS23 ZN C3 CX I3 C5 FP LA H21 LG NC HU2 RF C8 CN C7 I5 I8 I7
perturbations
D
### proteins
80
20>
cluster
1
2
3
4
log2 FC
>2
1
−1
−2>
not regulated
H2O1 HU AF DI FA I3 KA PBS C5 H21 CX NG HS42 ZN CS23 NC FP C8 LA C3 LG RF C7 CN I5 I8 I7
cluster
perturbations
E
cellular amino acid metab.
cytoplasmic translation
nuclear transport
nucleobase−containing metab.
monocarboxylic acid metab.
rRNA processing
cellular ion homeostasis
response to oxidative stress
ribosomal large sub. biogenesis
protein folding
carbohydrate metab.
cluster
−log10 q-value 1.3 2 3
enrichment
2 4 6 8
F
Glo3 S389
Sch9 S726
Elo2 T334
Sky1 S427
Nap1 T20
Sky1 S445
Hog1 Y176
Nsg2 S93
YPR091C T624
Rpc53 T232
Vps74 S14
Shp1 S106
Pil1 S45
Nup2 S205
Mrc1 S607
Hsp82 T612
Slt2 T190
Sla1 S996
Sic1 S201
Mck1 Y199
Fmp42 S249
Cho1 S42
Sky1 S388
Myo5 S777
Lhp1 S19
YJR056C S221
Rpa49 S151
Bim1 S165
YPR091C S716
YPR091C S714
Vps74 S19
Rpd3 S388
Pil1 S163
New1 T1191
log2 fold change vs untreated
### regulated perturbations
S389A
S726A
T334A
S427A
T20A
S445A
Y176A
S93A
T624A
T232A
S14A
S106A
S45A
S205A
S607A
T612A
T190A
S996A
S201A
Y199A
S249A
S42A
S388A
S777A
S19A
S221A
S151A
S165A
S716A
S714A
S19A
S388A
S163A
T1191A
### growth phenotypes p-site mutants (Viéitez et al)
ARFGAP2 432
RPS6KB1 403 / SGK1 401
ELOVL5 281
NAP1L4 12
MAPK12 185 / MAPK11 182 …
NLK 298 / MAPK7 219
SLC43A2 274
MYO1E 736 / MYO1F 734
MAPRE2 228 / MAPRE3 162
GOLPH3 36
conserved p-site in humans

**Figure S3 – Differential expression of phosphosites and proteins, related to Figures 1 and 3**

**(A)** Volcano plot displaying the negative $\log_{10}$ of the Benjamini-Hochberg corrected p-values by the $\log_2$ fold change determined by LIMMA for each phosphosite in each perturbation. **(B)** Same as (A) but for protein abundances. **(C)** Number of significantly regulated proteins in each selected perturbation. We quantified 2,185 proteins on average in each sample and found 257 regulated proteins and 791 regulated perturbation-protein pairs. Most perturbations showed regulation of less than 3% of all measured proteins. Intracellular pH changes had the strongest impact, affecting 5%-7% of measured proteins. **(D)** Heatmap and hierarchical clustering of regulated proteins across different perturbations. **(E)** Visualization of significantly enriched biological process terms within the 4 annotated clusters from (D). Most of the 123 proteins that increased in abundance showed a strong enrichment for amino acid metabolic processes, likely as a response to pH changes or adjusted metabolism. Down-regulated proteins were enriched for ribosome biogenesis, translation, and the oxidative stress response, all indicative of the early onset of the environmental stress response gene expression program. **(F)** Left: Dotplot of $\log_2$ fold changes of perturbations versus untreated for indicated phosphosites. Data points are color coded according to perturbation type (as in Figure 1) if the indicated phosphosite is significantly regulated. Middle: count of perturbations where the phosphosite is regulated. Right: count of significant phenotypes when exposing indicated phospho-inhibitory mutants to different stresses (Viéitez et al., 2022). Conserved phosphosites in human homologs are listed on the right.

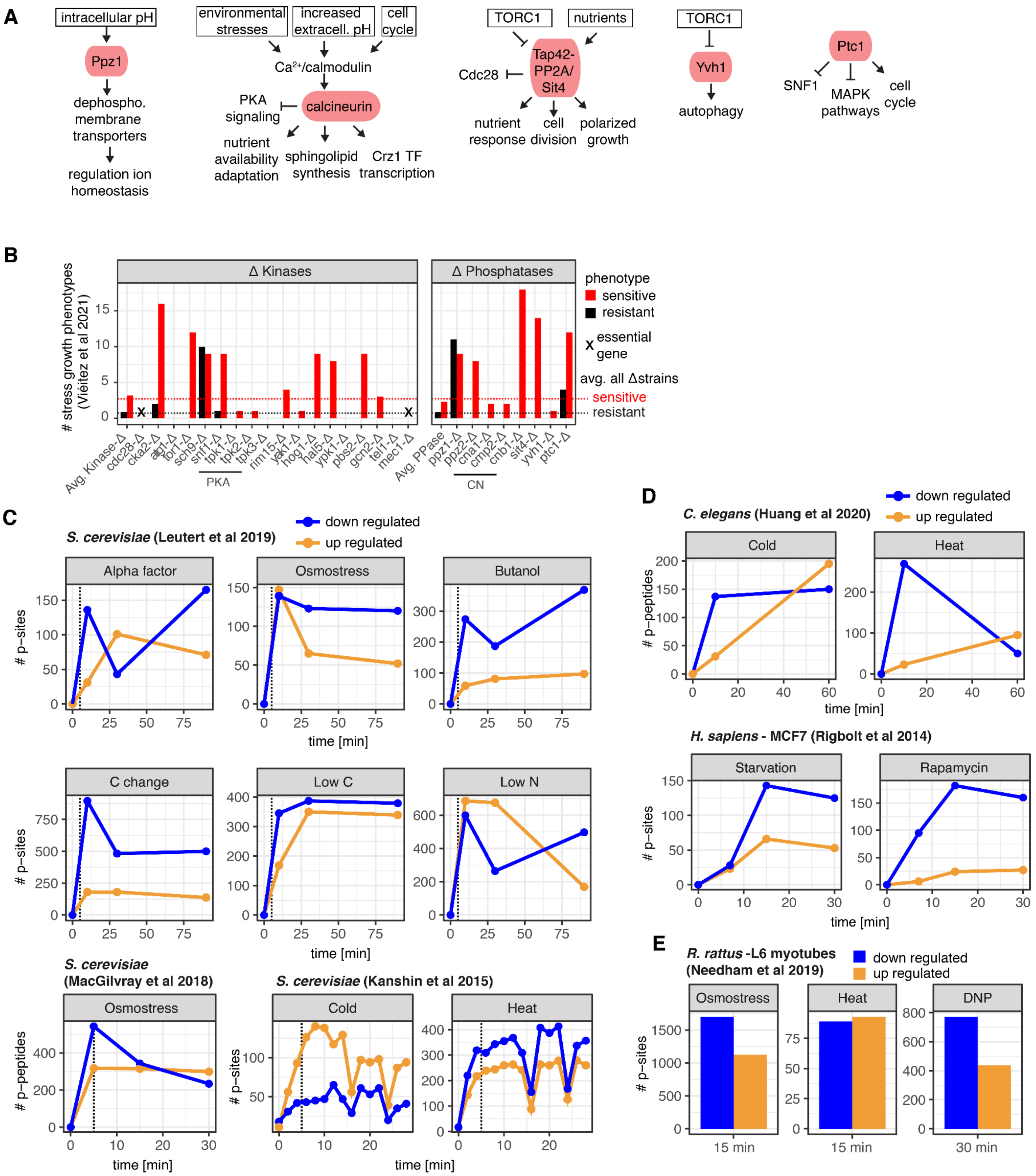
A
intracellular pH
Ppz1
dephospho. membrane transporters
regulation ion homeostasis
environmental stresses
increased extracell. pH
cell cycle
Ca²⁺/calmodulin
PKA signaling
calcineurin
nutrient availability adaptation
sphingolipid synthesis
Crz1 TF transcription
TORC1
nutrients
Cdc28
Tap42-PP2A/Sit4
nutrient response
cell division
polarized growth
TORC1
Yvh1
autophagy
Ptc1
SNF1
MAPK pathways
cell cycle
B
### stress growth phenotypes (Viéitez et al 2021)
Δ Kinases
Δ Phosphatases
phenotype
sensitive
resistant
X essential gene
avg. all Δstrains
sensitive
resistant
Avg. Kinase-Δ
cdc28-Δ
cka2-Δ
atg1-Δ
tor1-Δ
sch9-Δ
snf1-Δ
tpk1-Δ
tpk2-Δ
tpk3-Δ
rim15-Δ
yak1-Δ
hog1-Δ
hal5-Δ
ypk1-Δ
pbs2-Δ
gcn2-Δ
tel1-Δ
mec1-Δ
PKA
Avg. PPase
ppz1-Δ
ppz2-Δ
cna1-Δ
cmp2-Δ
cnb1-Δ
sit4-Δ
yvh1-Δ
ptc1-Δ
CN
C
S. cerevisiae (Leutert et al 2019)
down regulated
up regulated
### p-sites
Alpha factor
Osmostress
Butanol
C change
Low C
Low N
time [min]
S. cerevisiae (MacGilvray et al 2018)
Osmostress
### p-peptides
S. cerevisiae (Kanshin et al 2015)
Cold
Heat
### p-sites
time [min]
D
C. elegans (Huang et al 2020)
down regulated
up regulated
### p-peptides
Cold
Heat
time [min]
H. sapiens - MCF7 (Rigbolt et al 2014)
### p-sites
Starvation
Rapamycin
time [min]
E
R. rattus -L6 myotubes (Needham et al 2019)
down regulated
up regulated
### p-sites
Osmostress
Heat
DNP
15 min
15 min
30 min

**Figure S4 – Dephosphorylation is a major stress response, related to Figure 3**

**(A)** Summary of known regulation and functions of selected phosphatases (Ariño et al., 2019). **(B)** Count of stress-resistant and stress-sensitive growth phenotypes for deletion strains of selected kinases and phosphatases (indicated in Figure 4C) as determined by (Viéitez et al., 2022). Average phenotypes for all assessed kinase and phosphatase deletion strains are indicated by dashed lines. **(C)** Line plots of counts of down- and up-regulated phosphosites or phosphopeptides upon different perturbations of *S. cerevisiae* over time as identified previously (Kanshin et al., 2015; Leutert et al., 2019; MacGilvray et al., 2018). **(D)** Same plot as (C) for different perturbations in *C. elegans* and in the human MCF7 epithelial breast cancer cell line as previously identified by (Huang et al., 2020; Rigbolt et al., 2014). **(E)** Bar plot of down- and up-regulated phosphosite counts upon different perturbations in rat L7 myotubes as previously identified (Needham et al., 2019).

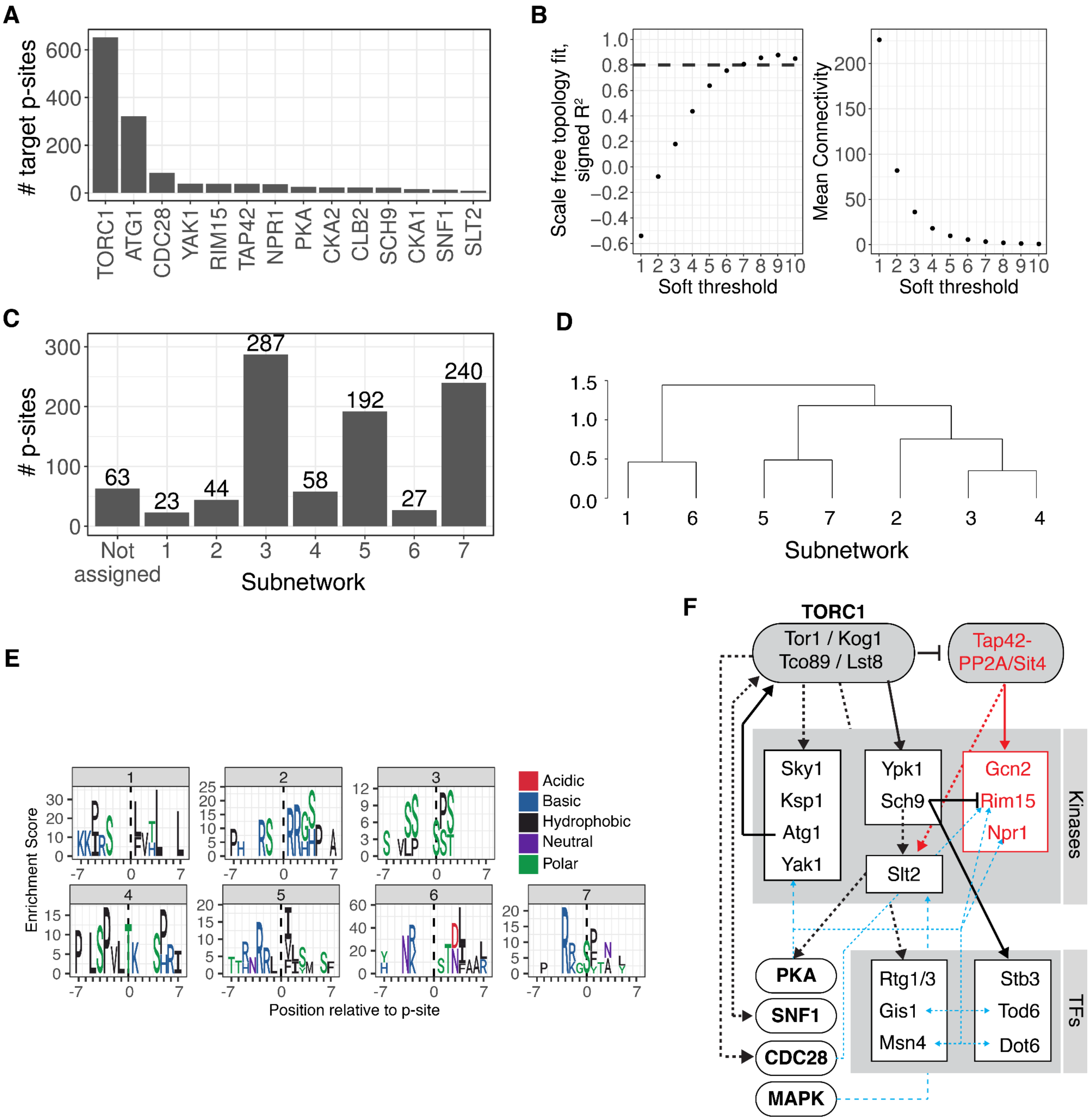


**Figure S5 – Analysis of the TOR signaling network, related to Figure 6**

**(A)** Numbers of target phosphosites associated with kinases in the TOR cascade that we considered in our analysis. **(B)** Signed $R^2$ of the scale free topology model fit (left) and the mean connectivity (right) of the underlying adjacency matrix produced by raising the Pearson correlation matrix to a soft threshold power. **(C)** Number of phosphosites in the TOR cascade assigned to each subnetwork discovered by WGCNA. **(D)** Hierarchical clustering of the Pearson correlation between module eigensites. **(E)** Linear motif enrichment of phosphosites assigned to each TOR subnetwork. **(F)** Schematic view of phosphoregulation of kinases and transcription factors associated with TOR signaling together with reported intersection points with other pathways.

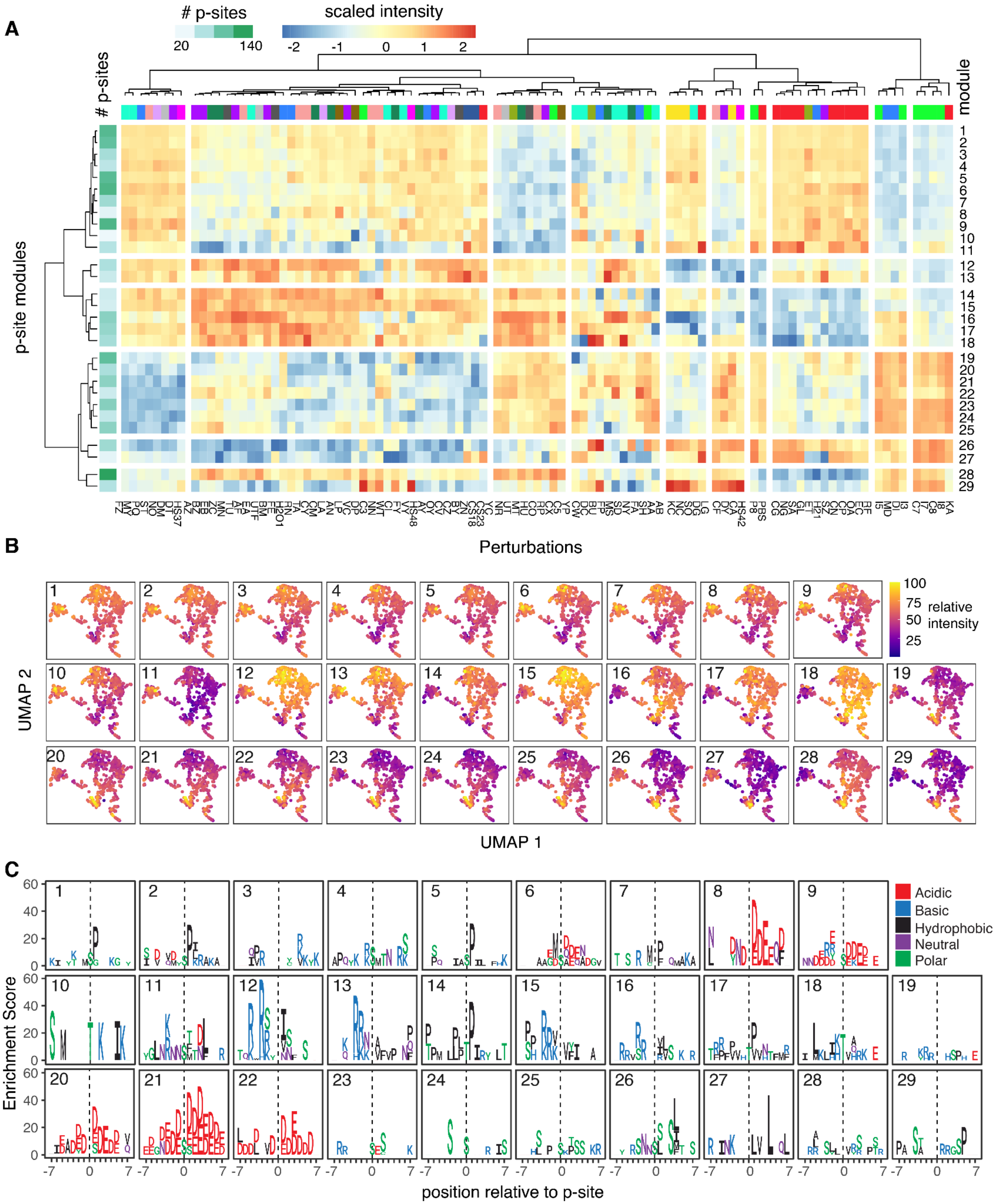
A
### p-sites
20
140
scaled intensity
-2
-1
0
1
2
### p-sites
module
p-site modules
Perturbations
B
UMAP 2
UMAP 1
100
75
50
25
relative intensity
C
Enrichment Score
position relative to p-site
Acidic
Basic
Hydrophobic
Neutral
Polar

**Figure S6 - Co-regulation Analysis, related to Figure 7**

**(A)** Heatmap showing the aggregated relative intensity of phosphosites within modules. The intensity is scaled across individual perturbations. Hierarchical clustering is performed on rows and columns. Perturbation types and phosphosites contained within a module are color-coded. **(B)** Aggregated relative intensity of all phosphosites within a module across the sample UMAP faceted for all modules. **(C)** Phosphosite motif enrichment analysis for all phosphosites within a module using all phosphosites assigned to modules as a background.

#### Supplementary Text

##### Overview

1. Dissecting stress responsive phosphorylation sites. Related to Figure 3D-F and Figure S3F.
2. Activity and architecture of the TOR signaling network. Related to Figure 6 and Figure S5.
3. Functional organization of the stress responsive phosphoproteome. Related to Figure 7 and Figure S6

##### 1. Dissecting stress responsive phosphorylation sites

Here we expand the analysis of individual phosphosites with well-characterized functions and their regulation across tested perturbations as reported in Figure 3D and Figure S3F. We discuss the regulation of these phosphosites in light of their function and pathway association.

We integrated regulation of functional sites within the cAMP-dependent protein kinase (PKA) pathway. PKA signaling plays a role in the regulation of cell growth, metabolism, and stress resistance and is constitutively active under optimal growth conditions. Neutral trehalase Nth1 S60 and S83 phosphorylations are important for Nth1 trehalose degrading activity (Dengler et al., 2021). Nth1 S60 and S83 phosphorylations were down-regulated in osmotic stresses, 42°C heat shock, and several nutrient and other perturbations (Figure 3D). Down-regulation under these perturbations suggests a build-up of trehalose, which functions as chemical stress protectant, and is required during osmotic stress, heat shock, and starvation. Atg1 S508, another PKA target, inhibits autophagy if nutrients are abundant (Budovskaya et al., 2005) and we found it was down-regulated upon carbon source perturbations (Figure 3D).

TORC1 is a major signaling hub that integrates diverse environmental cues and controls metabolic and biosynthetic processes. TORC1 is active if nutrients are abundant, inhibited following nutrient limitations, and in both cases unlocks various distally-controlled kinases and phosphatases to mediate cellular responses (González and Hall, 2017). TORC1-mediated phosphorylation of Sch9 kinase (ortholog of mammalian S6 kinase) is required for ribosome biogenesis, translation initiation, and entry into G0 phase (Urban et al., 2007). We observe strong down-regulation of activating Sch9 T723/T726 phosphosites upon nutrient and low pH perturbation (Figure 3D). Similarly, highly-conserved growth-promoting phosphorylations of S232/S233 on the ribosomal protein Rsp6a are down-regulated in nutrient perturbations, but also in many other perturbations, indicating a potential convergence of multiple TOR signaling axes and/or other pathways on Rps6a (Figure 3D). TORC1 inactivation promotes dephosphorylation of the effector Npr1 kinase at S51 and S317 leading to its activation. Npr1 regulates the activity and stability of several nutrient permeases at the plasma membrane (Dokládal et al., 2021). We observe broad dephosphorylation of these two phosphosites across nutrient perturbations (Figure 3D). Highly-conserved TORC1-dependent phosphorylation of superoxide dismutase Sod1 at S39 negatively regulates Sod1 enzymatic activity towards detoxifying reactive oxygen species. It is however not clear how TORC1-Sod1 redox regulation is induced (Tsang et al., 2018). We detect activation of Sod1 by dephosphorylation in increased pH perturbations, oxidative stress, and osmotic stress, but not in nutrient limitations, indicating a different mode of TORC1-dependent Sod1 phosphorylation compared to other analyzed TORC1 targets.

The SNF1 (AMP-activated protein kinase, AMPK) pathway is activated by carbon source depletion and essential for regulating a large set of genes and the activity of metabolic enzymes. SNF1 is also responsive to changes in nitrogen, pH, and salt levels and is involved in multiple stress responses. In agreement with its known functions, we observe strong up-regulation of known functional Snf1 targets involved in negative regulation of glucose-repressible genes (Reg1 S346), vesicle organization (Glo3 S389) and glycerol synthesis inactivation (Gpd2 S72) in nutrient limitations, but also in pH increase, and multiple other perturbations (Figure 3D). These findings establish SNF1 as another highly stress responsive signaling node.

Activating phosphorylation of the plasma membrane H+-ATPases Pma1, which is involved in pH homeostasis (Lecchi et al., 2007), was down-regulated in pH perturbations but also in nutrient limitations. Activating phosphorylation on heat shock transcription factor Hsf1 (Zheng et al., 2016) was induced during heat shock but also in certain oxidative and DNA damage insults and was down-regulated in pH perturbations. Activating phosphorylation on heat shock and redox transcription factor Skn7 (Kanshin et al., 2015b; Raitt et al., 2000) was up-regulated during cold shock and nutrient limitations. DNA repair inducing phosphorylation of histone H2A Hta1 S129 was up-regulated upon DNA damage treatments, but was down-regulated in pH stresses (Figure 3D).

Extensive bidirectional regulation was found for the highly conserved phosphorylation of the pyruvate dehydrogenase complex (PDH) subunit Pda1 on S313. PDH acts as a central metabolic node that connects glycolysis and the tricarboxylic acid (TCA) cycle. Phosphorylation of Pda1 S313 inactivates the PDH complex and the phosphorylation-deficient S313A mutant exhibits increased flux through PDH and the TCA cycle during growth on glucose (Oliveira et al., 2012; Uhlinger et al., 1986). We observe down-regulation of Pda1 S313 phosphorylation upon increase of extracellular pH, but up-regulation upon increase of intracellular pH (Figure 3F). Consistent with these results, increased TCA activity has been observed upon shift to media with unfavorable pH (Heyland et al., 2009). We hypothesize that the observed up-regulation of Pda1 S313 phosphorylation in intracellular pH changes, treatment with FCCP (protonophore) and deoxycholate (detergent affecting cell permeability) could be a fast switch to inhibit flux through TCA cycle upon mitochondrial damage and/or disruption of the mitochondrial proton gradient.

Seemingly counterintuitive, carbon starvation and carbon source changes led to PDH inactivation. Initial PDH inactivation might indicate that cells have to enter quiescence or undergo metabolic remodeling before committing to cellular respiration, similar to processes occurring during the diauxic shift.

#### 2. Activity and architecture of the TOR signaling network

Here we provide a detailed analysis and interpretation of individual subnetworks of the TOR pathway identified by WCNA and reported in Figure 6 and Figure S5.

Subnetworks 1 and 6 contained few and mostly anti-correlated phosphosites compared to the other subnetworks. Both subnetworks were up-regulated in osmotic stresses, pH, and C/N perturbations, although to a different extent (Figure 6B). Subnetworks 1 and 6 are strongly enriched for Snf1 kinase complex interactors (Figure 6C). Phosphosites show features of the Snf1 consensus motif, with a hydrophobic residue at the +4 position and upstream basic residues (Mok et al., 2010). Overall, these findings point towards targets regulated by Snf1 at the intersection of the SNF1-TOR networks. The Snf1 kinase complex is a major energy sensor activated by limited glucose and it has been shown that activated Snf1 negatively regulates TORC1 assembly (Hughes Hallett et al., 2015), which matches its anti-correlated activity compared to core TORC1 targets observed in other subnetworks. Although closely related, subnetwork 1 and 6 showed slightly different kinase consensus motifs and differential activation by the same perturbations, suggesting differences in upstream signaling .

Subnetwork 2 was strongly activated by osmotic stress (Figure 6B) and enriched in translation initiation factors (Figure 6C), suggesting that these phosphosites might be part of the osmotic stress-induced translation inhibition response. It was not clear if these sites are direct targets of TOR kinases or another pathway that shares common effector targets.

Subnetwork 3 was enriched in direct interactors of Tor1 kinase but not in any other TOR downstream kinases (Figure 6C). Subnetwork 3 contained the highest phosphosite counts on Gcn2, Npr1 and Rim15 kinases (Figure 6D), which are regulated through TORC1 mediated repression of the Tap42-PP2A/Sit4 phosphatase (Figure S5E). Subnetwork 3 was not affected by pH perturbation, but down-regulated in osmotic and nutrient stress conditions. This suggests that the mechanism of TORC1 in osmoregulation might partially stem from signaling through the Tap42-PP2A/Sit4 phosphatase complex.

Subnetwork 4 was exclusively down-regulated in nutrient limitations and its activity profile is most similar to subnetwork 3. Subnetwork 4 appeared to contain effector proteins further downstream of the TOR cascade, with a role in regulation of transport and response to chemicals (Figure 6C). Kinase interaction enrichment further suggests that subnetwork 4 might intersect with cell cycle control through Cdc28 (Figure 6C).

Phosphosites within subnetwork 5 showed a distinct basophilic motif with a hydrophobic (I, L, F) residue at the +1 position, which is indicative of direct TORC1 activity (Hsu et al., 2011; Hu et al., 2019; Yu et al., 2011) (Figure S5F). Phosphoproteins within subnetwork 5 were enriched in Tor1, Sch9 and Rim15 interactors. Subnetwork 5 contained TORC1-mediated activating phosphosites on Sch9 kinase and many additional phosphosites on kinases directly downstream of TORC1 and Sch9 (Figure 6D). Subnetwork 5 therefore best resembles the TORC1-Sch9 signaling branch. Subnetwork 5 was not only down-regulated in nutrient perturbations, but also in pH, osmotic, and other perturbations, representative of TORC1 function as a signal integrator for multiple stress types through TORC1-Sch9 (Figure 6B).

Subnetwork 7 is exclusively down-regulated in pH perturbations (Figure 6B) and shows similar activity profile and protein targets to subnetwork 5. Both subnetwork 5 and 7 contain phosphorylated interactors of PKA kinases. Subnetwork 7 contains many proteins directly in the TOR kinases cascade and downstream transcription factors (Figure 6C and 6D), including known activating phosphosphosites on Yak1 and Ksp1 kinases, which are directly targeted by PKA. Phosphosites in subnetwork 7 matched a basophilic motif with two arginines enriched at the -3 and -4 positions indicative of PKA or Sch9 activity (Plank et al., 2020). PKA and TORC1-Sch9 signaling are known to positively crosstalk and show a high level of interconnectivity at the level of downstream substrate phosphorylation with possible additive effects (Plank, 2022). TORC1-Sch9 and PKA signaling crosstalk has been previously proposed to function in pH homeostasis (Deprez et al., 2018)

#### 3. Functional organization of the stress responsive phosphoproteome

Here we provide detailed analysis and interpretation of selected modules of co-regulated phosphosites identified by the dimensionality reduction analysis and presented in Figure 7, Figure S6 and Table S9.

Module 12 contained 67 phosphosites on 55 proteins, included significantly down-regulated phosphosites in all nutrient perturbations, osmotic stresses, and external pH changes, showed a strongly basic phosphosite motif, and high enrichment for TORC1 and Sch9 kinase-protein interactors (Figure 7E). Phosphoproteins of module 12 included 9 kinases and 6 transcription factors.

Module 21 was down-regulated in heat and cold shock, as well as in perturbations targeting the cell cycle and cellular structure (Figure 7E). Both the acidic phosphosite motif and kinase-substrate enrichment analysis suggested casein kinase 2-dependent phosphorylation. Phosphoproteins were strongly enriched in nuclear localization and involved in ribosome biogenesis, cell cycle, and DNA metabolic process. Overall, these analyses indicate that module 21 might be involved in mediating cellular growth through casein kinase 2.

Module 5 was up-regulated in osmotic stress and heat shock, whereas module 17 shows the opposite regulation. Both modules show a proline-directed consensus motif. Module 5 is enriched in MAPK signaling pathway members, including 4 MAPK pathway associated kinases and 7 kinases not previously connected, as well as proteins involved in cytoskeleton organization. This suggests that phosphosites in module 5 are involved in, and targets of, MAPK signaling, which is the canonical response to high osmolarity and heat shock. In contrast, module 17 contained mostly cell cycle related proteins, as well as cellular budding, mitotic Gin4 kinase, and TOR pathway-associated proteins. The consensus motif and regulated targets indicate that co-regulation in this module is due to down-regulated activity of the master cell cycle kinase Ccd28.

Module 11 was up-regulated in nutrient limitations, osmotic and pH perturbations; and enriched consensus motif, kinase-protein interactions suggested activity of the canonical SNF1 pathway, affecting 60 phosphosphosites (Figure 7F).

Module 26 contained phosphosphosites that were up-regulated in carbon source perturbations, osmotic stresses, and high pH, and the phosphosite motif matches well with Kin1/Kin2 kinases (orthologous to MARK/PAR-1, AMPK family members). Phosphoproteins are enriched for Kin1 kinase protein interactors, plasma membrane proteins and regulation of transport (Figure 7E).
